# The Ubiquitin Ligase RBX2/SAG Regulates Mitochondrial Ubiquitination and Mitophagy

**DOI:** 10.1101/2024.02.24.581168

**Authors:** Wenjuan Wang, Ermin Li, Jianqiu Zou, Chen Qu, Juan Ayala, Yuan Wen, Md Sadikul Islam, Neal L. Weintraub, David J. Fulton, Qiangrong Liang, Jiliang Zhou, Jinbao Liu, Jie Li, Yi Sun, Huabo Su

## Abstract

Clearance of damaged mitochondria via mitophagy is crucial for cellular homeostasis. While the role of ubiquitin (Ub) ligase PARKIN in mitophagy has been extensively studied, increasing evidence suggests the existence of PARKIN-independent mitophagy in highly metabolically active organs such as the heart. Here, we identify a crucial role for Cullin-RING Ub ligase 5 (CRL5) in basal mitochondrial turnover in cardiomyocytes. CRL5 is a multi-subunit Ub ligase comprised by the catalytic RING box protein RBX2 (also known as SAG), scaffold protein Cullin 5 (CUL5), and a substrate-recognizing receptor. Analysis of the mitochondrial outer membrane-interacting proteome uncovered a robust association of CRLs with mitochondria. Subcellular fractionation, immunostaining, and immunogold electron microscopy established that RBX2 and Cul5, two core components of CRL5, localizes to mitochondria. Depletion of RBX2 inhibited mitochondrial ubiquitination and turnover, impaired mitochondrial membrane potential and respiration, and increased cell death in cardiomyocytes. *In vivo*, deletion of the *Rbx2* gene in adult mouse hearts suppressed mitophagic activity, provoked accumulation of damaged mitochondria in the myocardium, and disrupted myocardial metabolism, leading to rapid development of dilated cardiomyopathy and heart failure. Similarly, ablation of RBX2 in the developing heart resulted in dilated cardiomyopathy and heart failure. Notably, the action of RBX2 in mitochondria is not dependent on PARKIN, and PARKIN gene deletion had no impact on the onset and progression of cardiomyopathy in RBX2-deficient hearts. Furthermore, RBX2 controls the stability of PINK1 in mitochondria. Proteomics and biochemical analyses further revealed a global impact of RBX2 deficiency on the mitochondrial proteome and identified several mitochondrial proteins as its putative substrates. These findings identify RBX2-CRL5 as a mitochondrial Ub ligase that controls mitophagy under physiological conditions in a PARKIN-independent, PINK1-dependent manner, thereby regulating cardiac homeostasis.

## INTRODUCTION

The removal of damaged mitochondria through mitophagy, a selective form of autophagy, is critical for maintaining mitochondrial health and thus cellular hemostasis. Labeling damaged mitochondria with a phosphorylated ubiquitin (Ub) chain facilitates the recruitment of mitophagy receptors (i.e., p62, NBR1) and LC3-incorporated autophagosomes to eliminate dysfunctional mitochondria ^1–3^. This process is reportedly driven by two key proteins: the E3 Ub ligase PARKIN and the kinase PINK1 ^1, 2^. Following mitochondrial depolarization, PINK1 is stabilized and activated on the mitochondrial outer membrane (MOM), where it phosphorylates Ub at serine 65 (pS65-Ub) and activates PARKIN^4–6^. Active PARKIN, in turn, mediates the ubiquitination of other mitochondrial proteins for subsequent PINK1 phosphorylation, thereby resulting in a feed-forward loop and signal amplification.

Mitochondria play a crucial role in providing energy required for the continuously beating heart. In adult cardiomyocytes, mitochondria occupy approximately 30% of the cell volume^7^ and are constitutively turned over, as indicated by a half-life of around 6-17 days. ^8, 9^ Moreover, a high level of mitophagy activity has been detected in unstressed adult mouse hearts ^10, 11^. Nevertheless, PARKIN is expressed at very low levels in unstressed hearts ^12^, and germline or cardiac-specific deletion of *Parkin* had no impact on mitochondrial or cardiac function at baseline and during ageing.^13–15^ These findings suggest the existence of PARKIN-independent mitophagy in the heart. PARKIN-independent mechanisms have been linked to receptor-mediated mitophagy^16, 17^ and compensation via alternative Ub ligases such as HUWE1, ARIH1, MITOL and SIAH1^18–21^. However, germline deletion of *Bnip3* gene, or cardiac-specific ablation of *Bnip3l*, did not produce cardiac dysfunction ^22, 23^, arguing against a dominant role for receptor-mediated mitophagy in mitochondrial turnover in cardiomyocytes. While deletion of *Huwe1* and *Mitol* in the heart caused cardiomyopathy and heart failure^24, 25^, neither has been directly implicated in mitochondrial ubiquitination and turnover in cardiomyocytes. More recently, the ubiquitin ligase TRAF2 was shown to regulate mitophagy and cardiac homeostasis likely through a PARKIN-dependent mechanism^26^. Therefore, the ubiquitin ligases that control mitochondrial ubiquitination and clearance in cardiomyocytes under physiological conditions remain unidentified.

Cullin-RING Ub ligase 5 (CRL5) belongs to the Cullin-RING Ub ligase family, the largest family of Ub ligases ^27, 28^. CRL5 consists of four components: the scaffold protein Cul5, RING protein RBX2 (also known as SAG/ROC2/RNF7), adaptor proteins Elongin B/C and one of many substrate-recognizing receptors ^29^. Neddylation of cullins, which is catalyzed by NEDD8-specific E1-E2-E3 enzymes ^30^, is required for the assembly and activation of functional CRLs. As a core component of CRL5, RBX2 facilitates CUL5 neddylation and bridges Ub E2 to its substrates ^29^, which is essential for CRL5 activity. While RBX1, the other RING finger protein, shares 56% homology with RBX2, it primarily binds to other cullins (CULLIN 1, 2, 3, 4A and 4B) to promote their neddylation, leading to activation of CRLs 1-4. RBX2-CRL5 targets various substrates for proteolysis to regulate embryonic development, inflammation, viral infection, vasculogenesis and tumorigenesis ^31–37^. RBX2 is highly expressed in human and mouse heart ^38^. Global knockout of RBX2 in mice causes lethality at embryonic day 10.5 with pronounced cardiac edema ^36^. Despite these knowledges, the role of RBX2-CRL5 in mitochondrial turnover in the heart has not been explored.

In this study, we identify an association of CRLs with mitochondria. We show that RBX2 and Cul5, two core components of CRL5, localize to the mitochondria and are required for mitophagy in cardiomyocytes. Deletion of *Rbx2* in mouse hearts provokes accumulation of damaged mitochondria leading to the rapid development of heart failure and premature lethality. Mechanistically, loss of RBX2 inhibits mitochondrial ubiquitination and turnover and impairs mitochondrial respiration and function in cardiomyocytes in a PARKIN-independent, PINK1-dependent manner. Our results support a key role for the RBX2-CRL5 axis in maintaining mitochondrial and cardiac integrity mostly likely through directly ubiquitinating mitochondrial proteins.

## RESULTS

### Identification of CRL5 as a mitochondria-associated Ub ligase

We sought to identify Ub ligases other than PARKIN that are predominantly cytosolic but readily translocate to damaged mitochondria. Using peroxidase (APEX2)-mediated proximity biotinylation (Figure 1A), a recent study defined the mitochondrial outer membrane (MOM)-interacting proteome in human embryonic kidney (HEK) 293T cells ^39^. Cross referencing this MOM-interactome with the E3 Ub ligase database (ESI Network ^40^) identified 261 Ub ligase proteins that associate with mitochondria. Among these, 59 (∼22.6%) belong to the CRL family, including RBX1, RBX2, CUL1, CUL2, CUL3, CUL4B, CUL5, and their substrate receptors (Online Figure IA-B). Neddylation of Cullins induces the assembly of a functional multi-subunit CRL ^41^. Several neddylation enzymes were also identified as MOM-interacting proteins, including NEDD8 E1 enzyme NAE1 and UBA3, E2 enzyme UBE2M, and multiple subunits of deneddylase COP9 signalosome (CSN) (Online Figure IB). Similarly, overlap of the MOM-interactome with Ub ligases included in UbiNet 2.0 ^42^ confirmed the enrichment of CRLs in mitochondria (Online Figure IC).

**Figure 1.**
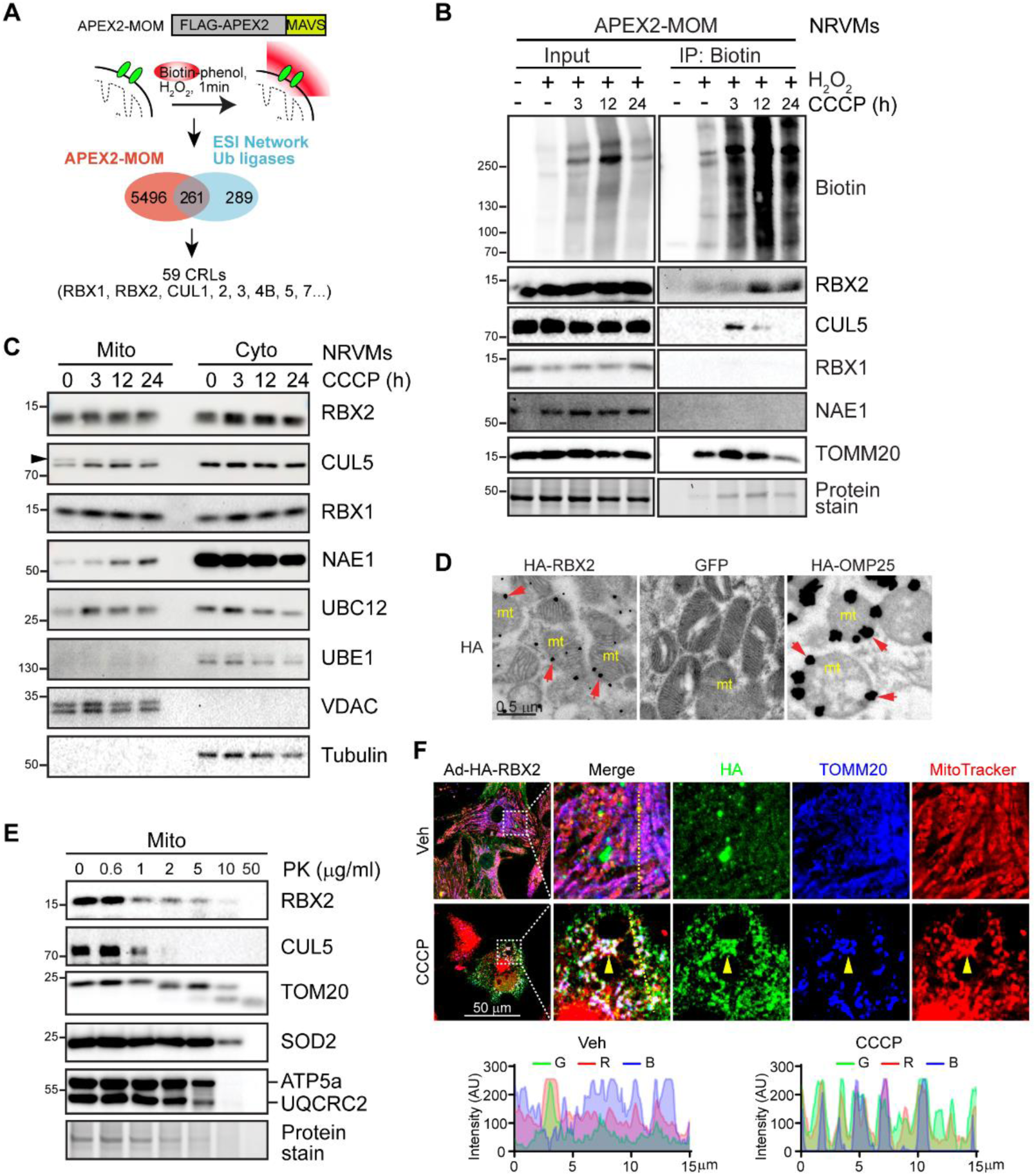
RBX2 and Cul5 localize to the mitochondria. **A**, Schematic of APEX2-catalyzed biotinylation at the mitochondrial outer membrane (MOM) via fusion to a mitochondrial targeted peptide derived from MAVS (mitochondrial antiviral-signaling protein). Overlap of APEX2-MOM data with the ESI (E3-substrate interaction) network reveals the association of CRLs with mitochondria. **B**, Representative Western blot of biotinylated proteins in neonatal rat ventricular cardiomyocytes (NRVCs) with adenoviral (Ad) expression of APEX2-MOM. NRVCs were treated with CCCP (10 µM) before H_2_O_2_ activation. The resultant biotinylated proteins were enriched by streptavidin beads. **C**, Representative Western blot of cytosol and mitochondrial fractions from NRVCs treated with CCCP (10 µM) for the indicated times. Arrowhead, neddylated CUL5. Tubulin and VDAC serve as cytosol and mitochondrial markers, respectively. **D**, Immuno-gold electron microscopic images showing the localization of RBX2 on mitochondrial membranes (arrowheads) in NRVCs expressing HA-RBX2. NRVCs infected with Ad-GFP and Ad-HA-OMP25 (MOM protein) serve as negative and positive controls, respectively. Scale bars, 0.5 µm. **E**, Representative Western blot of mitochondria with or without proteinase K (PK) treatment for 30 min on ice after isolation from NRVCs. **F**, Confocal images showing the colocalization (arrowhead) of RBX2 with mitochondria in CCCP-treated cardiomyocytes. NRVCs with adenoviral expression of HA-RBX2 were treated with CCCP for 3 hours and stained or immunostained as indicated. HA, green. Mitotracker, red. TOMM20, blue. Line scan co-localization analysis was done for all channels. Scale bars, 50 µm.

Mitochondria are abundant in cardiomyocytes. To validate the association of CRLs with mitochondria, neonatal rat ventricular cardiomyocytes (NRVCs) were infected with adenovirus expressing APEX2-MOM and treated with H_2_O_2_ to trigger biotinylation of MOM-interacting proteins. We detected the presence of MOM proteins (TOMM20, CISD1, TOMM40, SAMM50 and VDAC) and discernible levels of RBX2 in enriched biotinylated proteins at baseline (Figure 1B and Online Figure 1D). Interestingly, treatment with the mitochondrial uncoupler carbonyl cyanide chlorophenylhydrazone (CCCP) significantly increased the biotinylated forms of RBX2 and CUL5, but not those of RBX1, indicating enhanced association of RBX2 and CRL5 with mitochondria (Figure 1B). Consistently, Western blot following subcellular fractionation detected the presence of RBX2 and CUL5, as well as neddylation enzymes, in crude mitochondrial lysates (Figure 1C). Notably, neddylated CUL5 was predominant in mitochondria versus the cytosol, implying a role for active CRL5 in mitochondria. Immuno-gold electron microscopy detected the expression of exogenous RBX2 on mitochondrial membranes, consistent with the expression pattern of MOM protein OMP25 (Figure 1D). Moreover, in isolated mitochondria, RBX2 and CUL5, like the MOM protein TOMM20, were highly sensitive to proteolysis induced by proteinase K (PK), whereas mitochondrial inner membrane proteins ATP5a and UQCRC2, and matrix protein SOD2 (Figure 1E), were relatively resistant. Immunoreactive RBX2 was primarily detected in the cytosol and nucleus of untreated cardiomyocytes; however, CCCP treatment induced RBX2 recruitment to mitochondria labeled by MitoTracker and TOMM20 immunostaining (Figure 1F). Moreover, immunostaining of myocardium section also revealed the presence of RBX2 in mitochondria (Online Figure 1E). Together, these findings demonstrate that a subset of RBX2 and CUL5 proteins localize to the mitochondrial outer membrane.

### RBX2 mediates mitochondrial ubiquitination and turnover

To define the functional role of CRL5 in mitochondria, the catalytic subunit RBX2 was silenced in neonatal cardiomyocytes with two different siRNAs, which reduced CUL5 protein levels likely due to the disassembly and degradation of the CRL5 complex (Figure 2A and Online Figure IIA). Silencing of RBX2 markedly attenuated CCCP-induced pS65-Ub in both total cell lysates and mitochondrial fractions, in conjunction with reduced levels of P62 (autophagy receptor) and the accumulation of mitochondrial proteins (ATP5A, SDHB, MCQRC2, NDUFB8) (Figures 2A and 2B). Inhibition of proteasome function with Bortezomib (BZM) did not restore the levels of pS65-Ub and total ubiquitinated proteins in mitochondrial fractions in CCCP-treated, RBX2-deficient cardiomyocytes (Online Figure IIB), indicating that RBX2 does not target these mitochondrial proteins for proteasomal degradation. We next assessed mitolysosome numbers following incubation of cardiomyocytes with cell permeable Mtphagy dye ^43^, which revealed a significant reduction of mitophagic vesicles in RBX2-deficient cardiomyocytes at baseline and following CCCP treatment (Figure 2C and 2D). Moreover, the mitophagy flux assay showed a diminished turnover of LC3-II and P62 in mitochondria in RBX2-deficient cardiomyocytes, indicative of defective mitophagy (Figure 2E, Online Figure IIC). Together, these data demonstrate that RBX2 is necessary for mitochondrial ubiquitination and turnover in neonatal cardiomyocytes.

**Figure 2.**
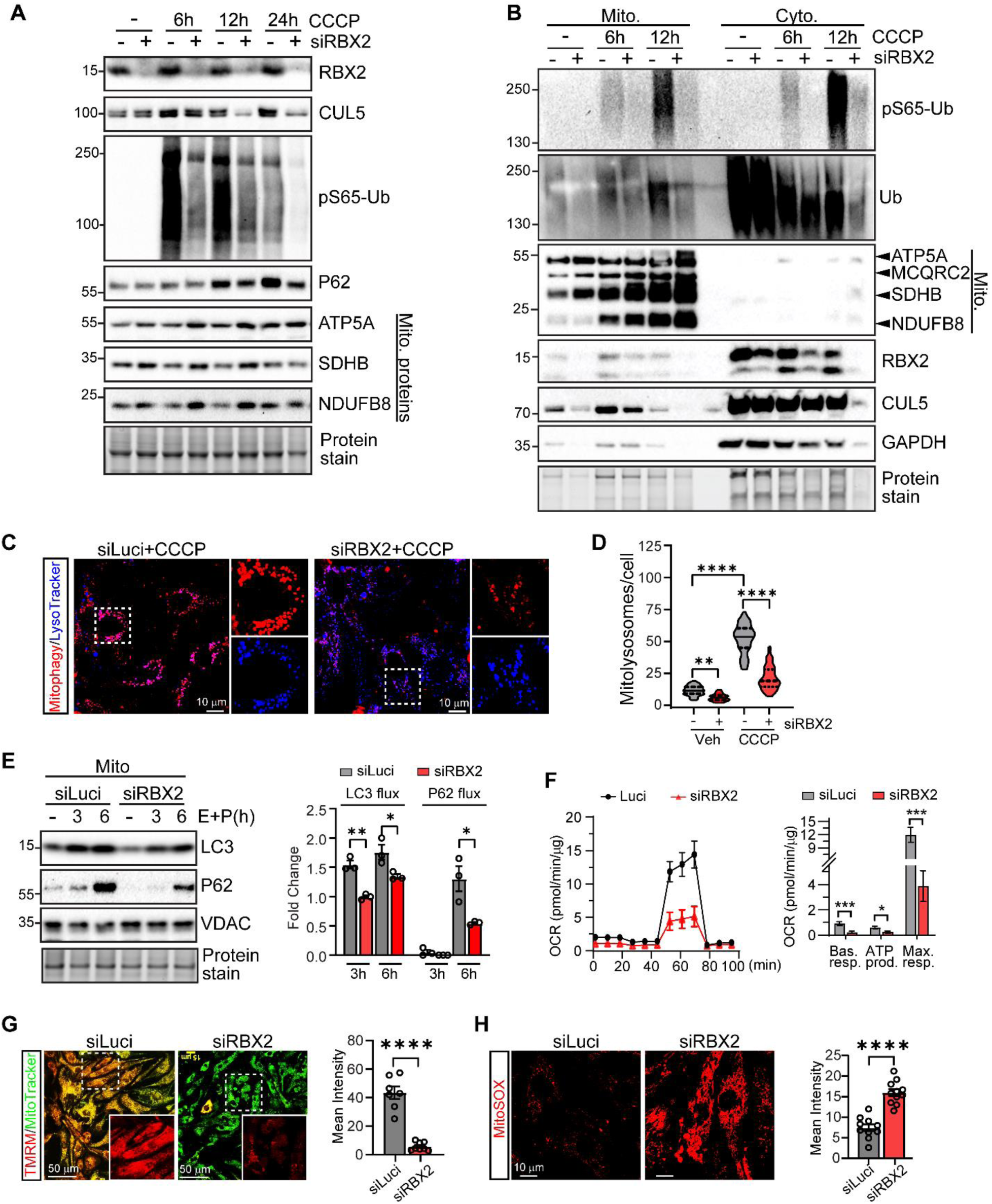
RBX2 mediates mitochondrial ubiquitination and mitophagy. **A**, Western blot of total cell lysates. NRVCs were transfected with indicated siRNAs, followed by CCCP (10 µM) treatment for the indicated times. **B,** Western blot of mitochondrial (mito) and cytosol (cyto) extracts. Cells were treated as described in **A**. **C,** Representative confocal images of live NRVCs stained with Mtphagy dye (red) and LysoTracker (blue) showing mitophagic vesicles. **D**, Quantification of mitophagic puncta per cell. **E,** Western blot of mitochondrial extracts. NRVCs were transfected with indicated siRNAs and treated with lysosomal protease inhibitors, E64d (E, 10 µg/ml) and pepstatin A (P, 10 µg/ml) for 6 hours before harvest. Fold-change of LC3-II and p62 levels between cells treated with and without lysosome inhibitors, indicative of mitophagy flux, is quantified on the right. **F**, Oxygen consumption rate (OCR) assessed by Seahorse analysis. Basal (Bas.) and maximal (Max.) respiration (resp.) are quantified on the right. siLuci: n=6, siRBX2: n=5 biological repeats. Data are representative of 3 repeats. **G**, Representative confocal images of NRVC stained with TMRM (1 µM, red) and MitoTracker (1 µM, green). The mean fluorescent intensity per view from 7-8 views (over 100 cells) per group is quantified on the right. **H**, Representative confocal images of NRVC stained with MitoSOX (red). The mean fluorescent intensity per view from 10 views (over 100 cells) per group is quantified on the right. One-way ANOVA was used in **D** and student *t* test in **E**-**H**. * *P* <0.05, ** *P*<0.01, *** *P*<0.001, **** *P* <0.0001.

Mitophagy maintains mitochondrial fitness and cellular health. Seahorse analysis demonstrated that depletion of RBX2 significantly decreased basal and maximal mitochondrial respiration and ATP production (Figure 2F). Furthermore, RBX2-deficient cardiomyocytes exhibited decreased mitochondrial membrane potential and elevated mitochondrial reactive oxygen species, as evidenced by TMRM and MitoSOX staining, respectively (Figure 2G-H). Consequently, RBX2-deficient cardiomyocytes were more susceptible to CCCP-induced cellular injury, as evidenced by augmented caspase 3 cleavage, elevated levels of lactate dehydrogenase release into medium, and increased cell death following CCCP treatment (Online Figure IID-IIF). Thus, these data suggest that RBX2 plays an important role in maintaining mitochondrial and cardiomyocyte integrity.

### RBX2 is indispensable for the function of adult mouse heart

We next generated tamoxifen (TAM)-inducible, cardiomyocyte-specific RBX2 knockout (RBX2^iCKO^, hereafter iCKO) mice by crossing *Rbx2^Flox^* mice with αMHC^MerCreMer^ (MCM) mice (Figure 3A). MCM mice exhibit transient and reversible cardiomyopathy following tamoxifen administration^44^ and are included as control. Tamoxifen administration (50 mg/kg/d for 5 days) induced efficient cardiac RBX2 depletion in 12-16-week-old iCKO mice compared with F/F and MCM mice. The iCKO hearts exhibited reduced levels of CUL5 protein, and modestly increased RBX1 protein, with no change in CUL3 or CUL4a (Figure 3B and 3C). Nearly 40% of iCKO mice died within two weeks after tamoxifen injection, while none of the F/F or MCM mice died during this period (Figure 3D). Compared with MCM and F/F mice, iCKO mice exhibited enlarged heart size and increased ratios of heart weight to tibial length and lung weight to tibial length (Figure 3E and 3F). Cardiac structure and function by echocardiography was indistinguishable between the three groups of mice prior to tamoxifen administration. However, iCKO mice promptly developed dilated cardiomyopathy and heart failure at 12 days after tamoxifen injections, as evidenced by left ventricular wall thinning, significant left ventricular chamber dilatation and severely impaired cardiac contractility (Figures 3G and 3H). Consistently, deletion of RBX2 led to pronounced cardiac pathology indicated by significantly increased cardiomyocyte cell size, marked cardiomyocyte apoptosis, upregulated expression of myocardial stress markers (*Acta1, Myh7*) and collagen (*Col1a*, *Col3a*), and reduced expression of *Myh6* (Figure 3I-K). To control for cardiotoxicity resulting from the MCM transgene^44^, we employed a two-week regimen of tamoxifen injections at lower doses (20 mg/kg/d for 10 days with a 2-day interval after the first 5 injections) (Online Figure IIIA). This regimen resulted in a significant reduction of RBX2 and CUL5 in Icko hearts (Online Figure IIIB) and led to cardiomyopathy (Online Figure IIIC-E). In contrast, MCM mice receiving this regimen of tamoxifen did not exhibit any alterations in myocardial RBX2 expression, heart weight, or cardiac functional/morphometric parameters as assessed by echocardiography (Online Figure IIIC-E). Cardiac dysfunction was not as pronounced following the low-dose tamoxifen injection protocol, possibly due to less efficient deletion of *Rbx2*. Collectively, these data demonstrate that RBX2 is necessary to maintain normal function of the adult heart.

**Figure 3.**
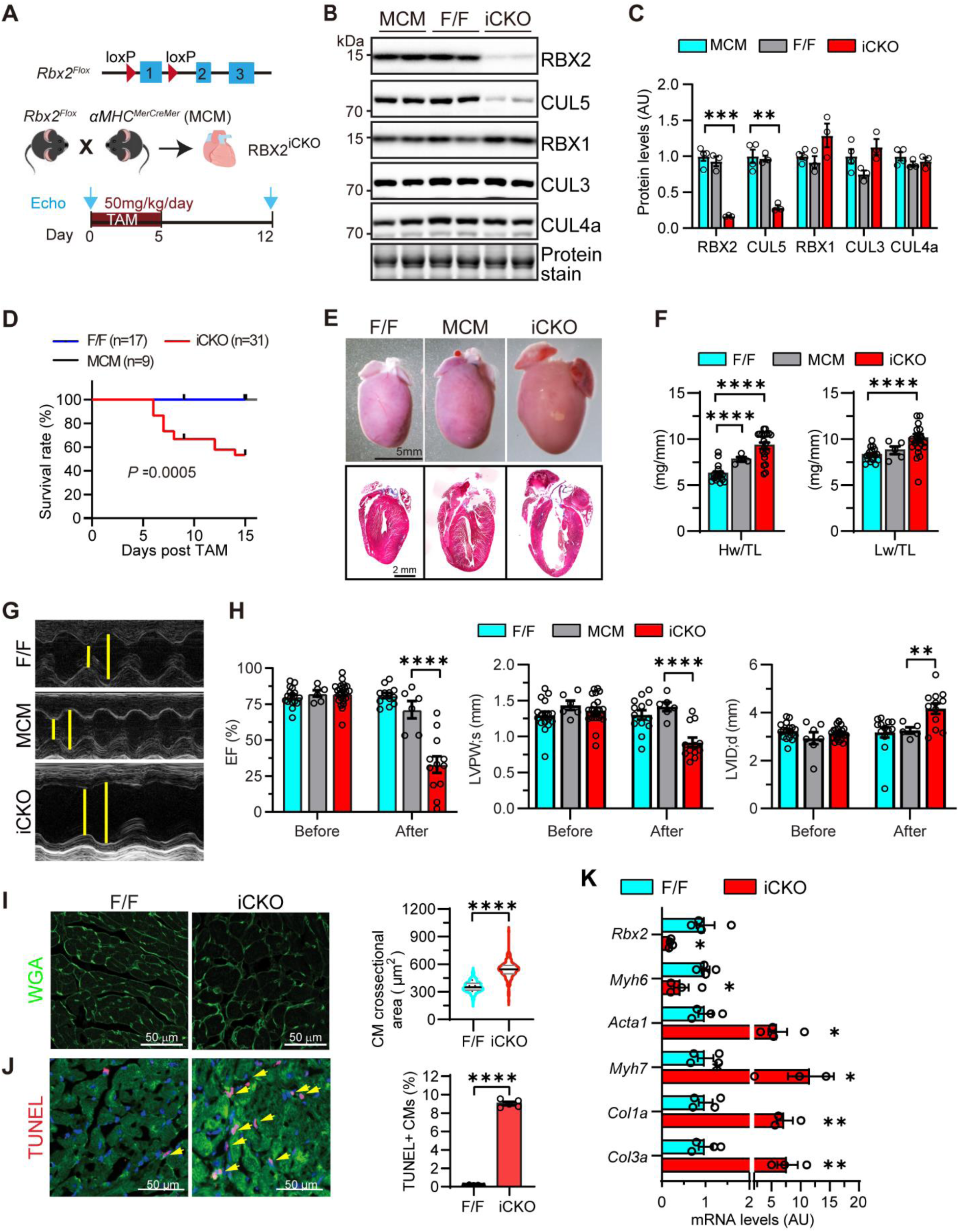
Deletion of *Rbx2* in adult heart leads to heart failure and lethality. **A**, Schematics of creation of tamoxifen-inducible, cardiac-specific RBX2 knockout (iCKO) mice. **B**, Western blot of indicated proteins in mouse hearts at 12 days after tamoxifen injection. **C**, Quantification of **B**. **D**, Survival curve. **E**, Gross morphology of mouse heart (top) and hematoxylin and eosin staining of myocardium section (bottom) at 12 days after tamoxifen injection. **F**, Heart weight to tibial length ratio and lung weight to tibial length ratio. F/F: n=13, MCM: n=5, iCKO: n=18. **G**, Representative B-mode images. **H**, Quantification of echocardiographic parameters before (F/F: n=17, MCM: n=6, iCKO: n=23) and after (F/F: n=12, MCM: n=6, iCKO: n=12) tamoxifen treatment. **I**, Wheat germ agglutinin (WGA) staining (left) of myocardium sections and quantification of cardiomyocyte (CM) cross-sectional area (right). More than 100 cells/heart and 3 and 12 hearts from F/F and iCKO mice, respectively, were quantified. **J**, Terminal deoxynucleotidyl transferase dUTP nick end labeling (TUNEL) staining of myocardium sections (left) and quantification (right). Three fields per heart, and 3 hearts per group, were quantified. **K**, qPCR analysis of the indicated genes. F/F: n=4, iCKO: n=4. One-way ANOVA followed by post hoc Tukey test was used in **C**, **F** and **H**. Log-rank (Mantel-Cox) test in **D**. Nested *t* test in **I** and **J**. Student *t* test in **K**. * *P* <0.05, ** *P*<0.01, *** *P*<0.001, **** *P* <0.0001.

### Perinatal deletion of RBX2 results in cardiomyopathy during ageing

RBX2 plays an important role in cell differentiation, proliferation and survival, and is essential for embryonic development in mice ^36, 37^. To determine the functional importance of RBX2 in the developing heart, we created cardiomyocyte-specific RBX2 knockout (RBX2^CKO^, hereafter CKO) mice by crossing *Rbx2^Flox^* mice with αMHC^Cre^ mice (Online Figure IVA). We confirmed significant reductions of RBX2 transcripts and proteins in RBX2^CKO^ hearts, which was accompanied by a robust reduction of CUL5 protein and an increase in the level of RBX1 protein, with little impact on its transcript levels (Online Figure IVB-D). While endothelial-specific deletion of RBX2 causes embryonic lethality due to defects in vasculogenesis^36^, CKO mice were viable and morphologically indistinguishable from their littermate controls. Serial echocardiography showed that compared with *Rbx2^F/F^* (F/F) mice and αMHC^Cre^ mice, RBX2^CKO^ mice exhibited discernible cardiac dysfunction (EF%: 65.9% of CKO vs 73.9% of *Rbx2^F/F^*) as early as 1 month of age and severe deterioration by 8 months of age (Online Figure IVE). As reported previously, αMHC^Cre^ mice develop cardiomyopathy during ageing^45, 46^. At 8 months of age, CKO hearts were enlarged compared with the hearts from F/F and αMHC^Cre^ mice (Online Figure IVE). Compared with 8-month-old F/F and αMHC^Cre^ mice, CKO mice exhibited more pronounced left ventricular wall thinning [LVPWs: 0.80 mm (CKO) vs 1.20 mm (αMHC^Cre^) and 1.32 mm (F/F)] and systolic dysfunction [EF%: 36.0% (CKO) vs 54.4% (αMHC^Cre^) and 80.2% (F/F)] (Online Figure IVE-H). Moreover, CKO hearts exhibited more myocardial interstitial fibrosis and heightened expression of cardiac stress maker genes such as *Nppa*, *Nppb*, *Mhy7 and Acta1*, and downregulation of *Atp2a2* expression, compared with littermate F/F hearts (Online Figure IVI-J), indicative of adverse cardiac remodeling. These data support a crucial role of RBX2 in maintenance of cardiac homeostasis.

### RBX2 deficiency has a major impact on the mitochondrial proteome

To gain mechanistic insight into how RBX2 regulates cardiomyocyte function, we sought to define the RBX2 regulated proteome. Given the likely confounding effects resulting from chronic RBX2 depletion *in vivo*, we performed tandem mass tag (TMT)-based quantitative proteomics analyses on total cell lysates from control (CTL) and RBX2-deficient (KD) cardiomyocytes in the absence and presence of CCCP treatment (Figure 4A). Principal component analysis (PCA) of protein abundances showed distinct separation between the different groups (Figure 4B). Mass spectrometry (MS) analysis detected 5199 proteins, of which 2425 were quantitatively detected in all 10 replicates from four groups. The results did not identify any differentially expressed proteins (DEPs) between CTL and KD groups, between CTL+CCCP and CTL groups, and between KD+CCCP and KD groups (FDR<0.1, FC>1.3 or <0.77) (Figure 4C). These data indicate that RBX2 deficiency had a negligible impact on the global proteome at baseline, and that CCCP treatment for 6 hours did not significantly alter the proteome in CTL and RBX2-deficient cardiomyocytes. In contrast, following CCCP treatment, ∼18.5% of the quantitively detected proteins were dysregulated (180 downregulated and 269 upregulated) in RBX2-deficient cardiomyocytes compared with CCCP-treated CTL cardiomyocytes (Figure 4C and 4D, Online Table 1). Gene ontology analysis identified the enrichment of upregulated proteins in the pathways relevant to oxidative phosphorylation (FDR < 2.2 x 10^^-51^), diabetic cardiomyopathy (FDR < 5.3 x 10^^-50^), cardiac muscle contraction (FDR < 1.1 x 10^^-14^), electron transportation chain (FDR < 1.6 x 10^^-17^) and in mitochondrial subcompartments such as matrix (FDR < 4.6 x 10^^-16^) and outer membrane (q < 2.1 x 10^^-8^), whereas the downregulated proteins are enriched in pathways related to actin cytoskeleton organization (FDR < 5.8 x 10^^-14^) and the sarcomere (FDR < 3.6 x 10^^-11^) (Online Figure VA-VB). Notably, crossing the dysregulated proteome with mitochondrial proteins annotated in MitoCarta 3.0 demonstrated that 127 (47.2%) of the 269 upregulated proteins are mitochondrial proteins (Figure 4E), whereas only 10 (5.6%) of the 180 downregulated proteins are in mitochondria. These dysregulated mitochondrial proteins localize in different mitochondrial subcompartments (Online Figure VC and VD). These data are consistent with a critical role for RBX2 in maintaining mitochondrial protein homeostasis and the regulation of mitochondrial turnover.

**Figure 4.**
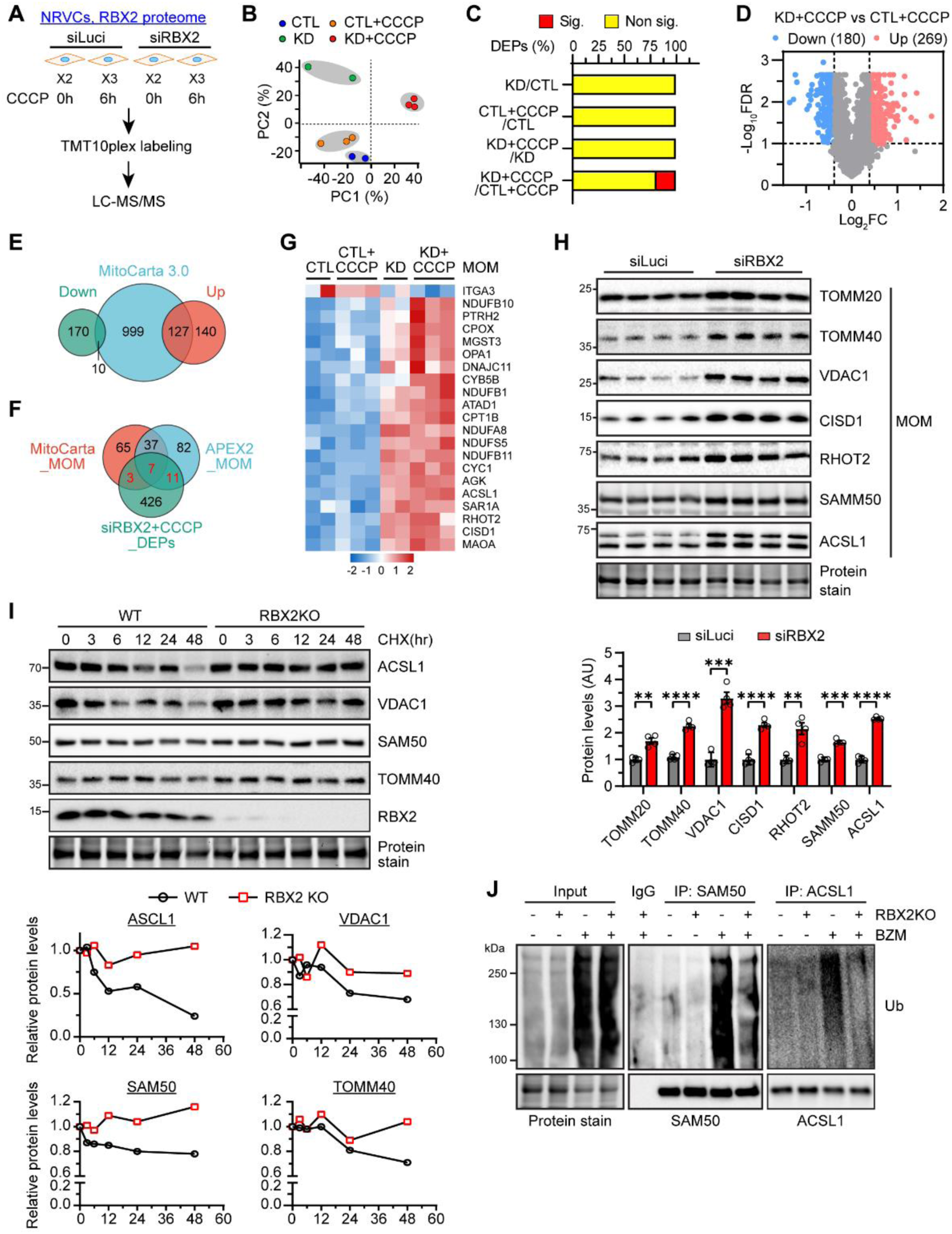
RBX2-regulated mitochondrial proteome. **A**, Scheme of procedures for identification Rbx2-regulated proteome in NRVCs. NRVCs were transfected with indicated siRNAs, followed by CCCP (10 µM) treatment for 6 hours. Cell lysates were collected for trypsin digestion. The resultant peptides were labeled by TMT before mass spectrometry analysis. **B**, Principal component analysis (PCA) of normalized protein expression in whole cell proteome. CTL, siLuci. KD, siRBX2. **C**, Analysis of the percentage of differential expression of proteins (DEPs) in each group. **D,** Volcano plot of differentially expressed proteins (Sig., blue and red) in CCCP-treated RBX2-deficient CMs (KD) compared with CCCP-treated CTL. **E**, Venn diagram showing overlap of RBX2-regulated proteome and mitochondrial proteins annotated in MitoCarta 3.0. **F**, Venn diagram showing the identification of RBX2-regulated MOMs. **G**, Heatmap showing the relative expression of MOM proteins amongst the groups of cells. **H**, Western blot of MOM proteins in control and RBX2-deficient cardiomyocytes. N=4 for each group. Multiple *t* test was used. ** *P*<0.01, *** *P*<0.001, and **** *P*<0.0001. **I**, Western blot of WT and RBX2KO Hela cells treated with cycloheximide (CHX, 100 nM) for indicated times. *RBX2* was deleted in Hela cells via CRISPR/Cas9-mediated gene editing. **J**, WT and RBX2KO Hela cells were transfected with plasmids expressing HA-Ub (pCDNA3-HA-Ub), treated with a proteasome inhibitor Bortezomib (BZM, 100 nM) for 6 hours, and subjected to immunoprecipitation followed by Western blot.

To explore which mitochondrial proteins are RBX2 substrates, we crossed the mitochondrial DEPs with MOMs annotated in MitoCarta 3.0 and the APEX2-MOM study ^39^, respectively. The results identified 21 MOMs as potential RBX2 substrates (10 from MitoCarta_MOM and 18 from APEX2_MOM) (Figure 4F). All of these MOMs except ITGA3 were upregulated in the KD+CCCP group compared with the CTL+CCCP group and showed a trend towards upregulation, though not statistically significant, in the KD group compared with the CTL group (Figure 4G). Furthermore, among 112 MOM proteins annotated in MitoCarta 3.0, 37 were detected by our assay and 10 were significantly upregulated in RBX2-deficient cardiomyocytes (Online Figure VE). Immunoblotting confirmed the upregulation of these MOM proteins (TOMM20, TOMM40, VDAC1, CISD1, RHOT2, SAMM50, ACSL1) and MIM proteins (HSP60, AGK, CYC1, NUDUFS5), without discernable changes in their transcript levels in RBX2-deficient cardiomyocytes (Figure 4H, Online Figure VF and VG). Cycloheximide-based pulse chase assays showed reduced degradation rate of ASCL1, VDAC, SAM50 and TOMM40 in RBX2-deficient Hela cells (Figure 4I). Moreover, immunoprecipitation showed that RBX2 deficiency inhibits the ubiquitination of SAM50 and ASCL1 (Figure 4J). Together, these data suggest that RBX2 controls the ubiquitination and degradation of a subset of MOMs.

### RBX2 deficiency inhibits mitochondrial ubiquitination and mitophagic activity in the heart

We next sought to examine the impact of RBX2 deficiency on cardiac mitochondrial turnover in iCKO hearts. MCM mice receiving tamoxifen exhibited metabolic dysfunction^44^ and were included as stringent controls. Consistent with a previous report^47^, detection of phosphorylated ubiquitin in tissues is challenging and we were not able to detect the myocardial levels of phosphorylated ubiquitin by western blot using total heart lysates, likely due to a relatively low percentage of mitochondria undergoing mitophagy in the normal heart at a given time point. Immunostaining of pUb on myocardium sections from MCM detected pUb-positive mitochondria, which were significantly reduced in iCKO hearts (Figure 5A and 5B). During mitophagy, ubiquitinated damaged mitochondria are recognized and delivered to autophagosomes for lysosomal degradation by adaptor proteins such as p62 ^48^. Western blot showed less mitochondrial P62, but not those in cytosol, in iCKO hearts compared with MCM hearts (Figure 5C and 5D). Consistently, immunostaining also detected diminished co-localization of p62 with mitochondria in iCKO hearts (Online Figure VIA). We further performed autophagy flux assay in mouse hearts by treating MCM and iCKO mice with a lysosome inhibitor bafilomycin A1 (BFA). Western blot showed that loss of RBX2 suppressed LC3-II turnover in mitochondrial fraction but had no effects on LC3-II turnover in cytosolic faction in mouse hearts (Figure 5E and 5F). Examination of mitophagic activity via AAV9-mediated transduction of mt-Keima reporter^49, 50^ demonstrated marked suppression of mitophagy in iCKO hearts (Figure 5G and 5H). Assessment of mitophagic activity in 2-month-old CKO hearts, in which cardiac function is largely maintained compared with littermate controls (Figure 3D), also demonstrated decreased mitophagy activity (Online Figure VIB and VIC). Furthermore, in isolated adult cardiomyocytes, RBX2 deficiency reduced the levels of phosphorylated ubiquitin (Figure 5I) and CCCP-induced mitophagic vesicles (Online Figure VID and VIE). Together, these *in vivo* and *ex vivo* data suggest that RBX2 is required for the ubiquitination of mitochondria proteins and mitochondrial turnover in the heart.

**Figure 5.**
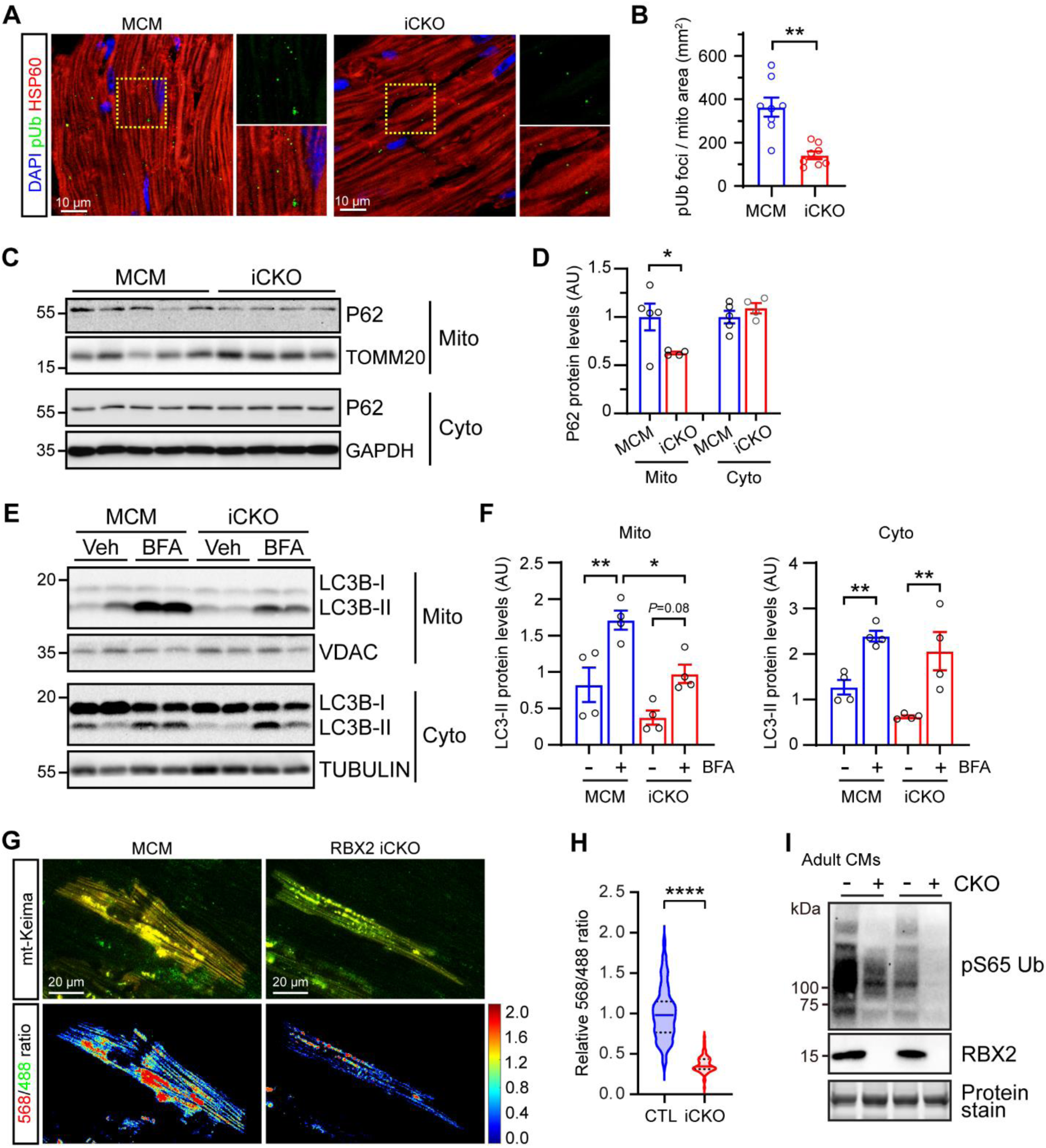
Impaired mitochondrial ubiquitination and mitophagy in RBX2-deficient hearts. Adult MCM and RBX2^iCKO^ (iCKO) mice were administered with tamoxifen (50 mg/kg/d for 5 days). Tissues were collected for indicated analyses (**A**-**H**) at 12 days after tamoxifen injections. **A**, Representative confocal images (left) of MCM and RBX2^iCKO^ myocardium sections immunostained with pUb (green), HSP60 (red, mitochondrial marker) and DAPI. Scale bars, 10 µm. **B**, Quantification of pUb+ foci normalized by mitochondria (HSP60+) area. A total of 16 views from two hearts per group were quantified. **C**-**D**, Western blot (**C**) and quantification (**D**) of mitochondrial (mito) and cytosolic (cyto) P62 in mouse hearts. **E**-**F**, Western blot (**E**) and quantification (**F**) of mitochondrial (mito) and cytosolic (cyto) LC3-II in mouse hearts. Mice at 12 days after tamoxifen administration were intraperitoneally injected with bafilomyocin A1 (BFA, 3 µmol/kg) for 3 hours before tissue harvest. **G**, Representative confocal images of mt-Keima at 488 nm and 568 nm, respectively, in epicardial cardiomyocytes and the derived heatmaps. Neonatal MCM and RBX2^iCKO^ mice were transduced with AAV9-mt-Keima (1X10^11^ GC/pup). At 10 weeks of age, mice were treated with tamoxifen and intact mouse hearts were excised 12 days later and scanned for mt-Keima signals in epicardial cardiomyocytes *in situ* with confocal microscope. Scale bars, 20 µm. **H**, Quantification of relative 568/488 ratio. 20-40 views per heart, 3 hearts per group were quantified. **I**, Western blot of indicated proteins in adult cardiomyocytes isolated from 2-month-old CTL (RBX2^F/F^) or RBX2^CKO^ (CKO) mouse hearts. Results from two different batches of cells are shown. Nested *t* test was was used in **B** and **H**, Mann-Whitney test in **D**, and One-way ANOVA in **F. \*** *P* < 0.05, **\*\*** *P* < 0.01, **** *P* <0.0001.

### Loss of RBX2 disrupts metabolic pathways and impairs mitochondrial homeostasis

To gain additional mechanistic insight into how RBX2 regulates cardiac function, we performed transcriptomic analyses of CTL (F/F) and iCKO hearts. Principal component analysis showed that CTL and mutant hearts exhibited distinct gene expression patterns (Online Figure VIIA). A total of 453 downregulated and 881 upregulated genes were identified in RBX2-deficient hearts (FC > or < 1.5, *P*_adj_ < 0.05. Online Figure VIIB). Kyoto encyclopedia of genes and genomes (KEGG) and gene ontology analyses demonstrated enrichment of downregulated genes in fatty acid degradation, PPAR signaling and cardiac muscle contraction, while upregulated genes were enriched in extracellular matrix remodeling, relaxin signaling and PI3K-Akt signaling, among others (Figure 6A, Online Figure VIIC). In particular, many downregulated genes encoded mitochondrial proteins that are involved in fatty acid oxidation, oxidative phosphorylation and the electron transport chain (Figure 6B, Online Figure VIID), suggesting deficits in oxidative metabolism and myocardial bioenergetics. Transmission electron microscopy demonstrated ultrastructural abnormalities in RBX2-deficient cardiomyocytes, including focal myofibrillar lysis, enlarged and swollen mitochondria with disrupted cristae associated with lysosomes, massive mitophagic vesicles, and dilated endoplasmic reticulum (ER) (Figure 6C). Immunostaining of myocardium sections revealed a substantial increase of LAMP1 (a lysosome marker)-positive mitochondria in iCKO hearts compared with MCM hearts (Figure 6D), indicative of accumulation of damaged mitochondria. Despite the diminished mitochondrial ubiquitination, the increased mitophagic vesicles in RBX2-deficient hearts suggest the possible contribution of additional Ub ligases and receptor-mediated mitophagy to mitochondrial clearance. Interestingly, the ubiquitin ligase TRAF2 was recently reported to mediate mitophagy in the heart^26^ and was upregulated in iCKO hearts, whereas expression of BNIP3 and BNIP3L, key regulators of receptor-mediated mitopohagy^16, 17^, was not affected (Online Figure VIIE). Seahorse analysis further confirmed compromised mitochondrial bioenergetics in RBX2-deficient hearts (Figure 6E). Consistent with the transcriptomics analysis, immunoblotting showed that loss of RBX2 resulted in substantial downregulation of mitochondrial proteins (ATP5A, UQCRC2, SDHB, NDUFB8) in the heart (Figure 6F). We speculate the transcriptomics changes could represent adaptive responses to mitochondrial stress resulting from defective mitochondrial turnover and/or concomitant pathological cardiac remodeling. Moreover, Parkin and PINK1 were upregulated and downregulated in RBX2-deficient hearts, respectively (Figure 6F). Collectively, these *in vivo* data, coupled with our *in vitro* findings, are consistent with the critical role for RBX2 in maintaining mitochondrial homeostasis and function in the heart.

**Figure 6.**
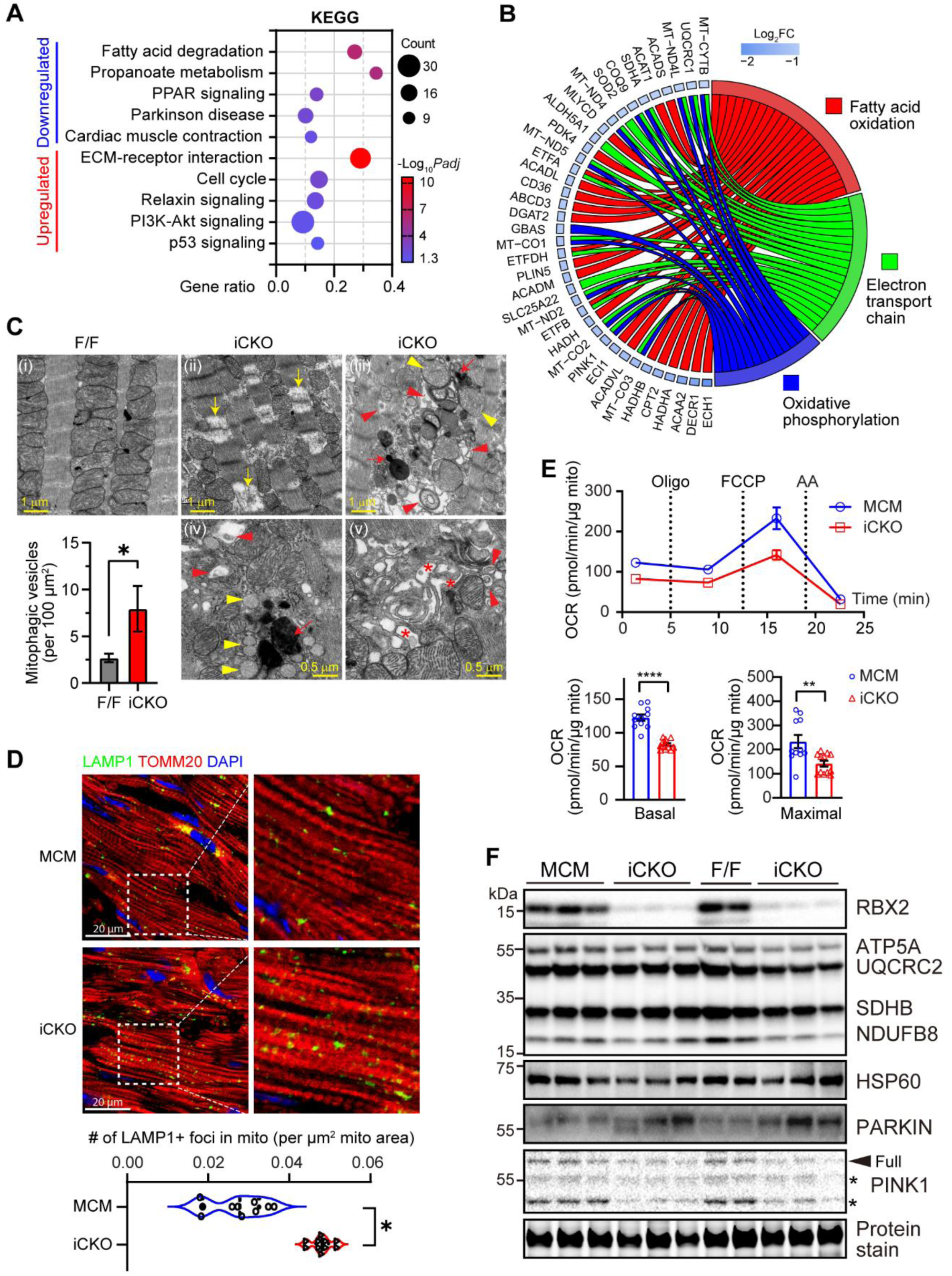
Alterations in metabolic pathways and mitochondrial homeostasis in RBX2-deficient hearts. **A**-**B**, Bulk RNA sequencing of mouse hearts at 12 days after tamoxifen (50 mg/kg/day for 5 days) treatment. **A**, KEGG pathways enriched in downregulated (top) and upregulated (bottom) genes in iCKO hearts. **B**, Chord plot showing downregulated genes involved in the indicated metabolic processes. **C**, Transmission electron microscopic images showing myofibril lysis (*), degenerating mitochondria (yellow arrowheads) surrounding by lysosomes (arrows), and abundant mitophagic vesicles (red arrowheads) in mutant cardiomyocytes. The number of mitophagic vesicles was quantified. Over 50 cardiomyocytes per group were quantified. **D**, Representative confocal images of myocardium sections immunostained with LAMP1 (green), TOMM20 (red) and DAPI (blue). Quantification of LAMP1+ puncta in mitochondria is shown. 5-6 views per heart, 2 hearts per group were quantified. **E**, Seahorse analysis of mitochondria isolated from mice at 8 days after tamoxifen injections. Basal and maximal oxygen consumption rates are shown. **F**, Western blot of mitochondrial proteins in mouse hearts. *, cleaved form. Nested *t* test was used in **C** and **D** and student t-test in **E**. * *P*<0.05. ** *P*<0.01, and **** *P*<0.0001.

### PARKIN is dispensable in RBX2-mediated mitophagy

Given the central role of PARKIN in mitochondrial ubiquitination, we asked whether RBX2 acts through PARKIN to mediate mitochondrial ubiquitination and mitophagy. Silencing of RBX2 did not alter *Parkin* transcriptional levels (Online Figure VIIIA). Since PARKIN protein expression was very low in neonatal cardiomyocytes, we probed the effect of RBX2 on PARKIN expression in cardiomyocytes infected with Ad-PARKIN. Our results showed that under these conditions, RBX2 depletion insignificantly reduced PARKIN protein levels (Figure 7A). Moreover, in SH-SY5Y cells, a PARKIN-expressing neuroblastoma cell line, deletion of *RBX2* did not alter PARKIN expression at baseline but resulted in accumulation of PARKIN protein following CCCP treatment (Online Figure VIIIB). Furthermore, we detected increased PARKIN protein levels in RBX2-deficient mouse hearts (Figure 6G). Thus, loss of RBX2 does not lead to reduced PARKIN expression. To further determine the contribution of PARKIN to RBX2-mediated mitochondrial ubiquitination, cardiomyocytes were isolated from neonatal wild-type and PARKIN null mice ^51^. Deletion of *Rbx2* exhibited similar inhibitory effects on CCCP-induced pS65-Ub in wild-type and PARKIN null cardiomyocytes and cardiac fibroblasts (Figure 7B, Online Figure VIIIC). Similarly, depletion of RBX2 exerted a greater repressive effect on pS65-Ub than depletion of PARKIN via siRNA (90% reduction in mRNA by qPCR, Online Figure VIIID) and remained effective in PARKIN-depleted cardiomyocytes (Figure 7C). Moreover, PARKIN overexpression dose-dependently promoted pS65-Ub following CCCP treatment, which was attenuated by siRNA-mediated depletion of RBX2 (Figure 7D), suggesting that RBX2 is required for PARKIN-mediated mitochondrial protein ubiquitination under these conditions. We compared the RBX2-regulated 137 mitochondrial proteins with 47 PARKIN mitochondrial substrates that were previously identified in HCT116 and Hela cells ^52^. We detected only 9 overlapping proteins; among these, 4 are MOM proteins (Online Figure VIIIE), suggesting that RBX2 and PARKIN have distinct mitochondrial substrates. These data suggest that RBX2 does not require PARKIN to mediate mitochondrial ubiquitination.

To examine potential interactions between RBX2 and PARKIN in the heart, we generated Parkin and RBX2 double knockout (DKO) mice by breeding PARKIN knockout (*Parkin^-/-^*) mice ^51^ with CKO mice (Figure 7E). Consistent with previous reports^13–15^, PARKIN deficiency did not impair cardiac contractility in 8-month-old mice (EF%, 77.3±4.4%). Importantly, deletion of *Parkin* in CKO mice did not alter the onset or progression of dilated cardiomyopathy as demonstrated by temporal echocardiography, nor did it affect the survival rate of the CKO mice (Figure 7F and 7G, Online Figure VIIIF). Together, these data suggest that RBX2 regulates cardiac homeostasis in PARKIN-independent mechanism.

**Figure 7.**
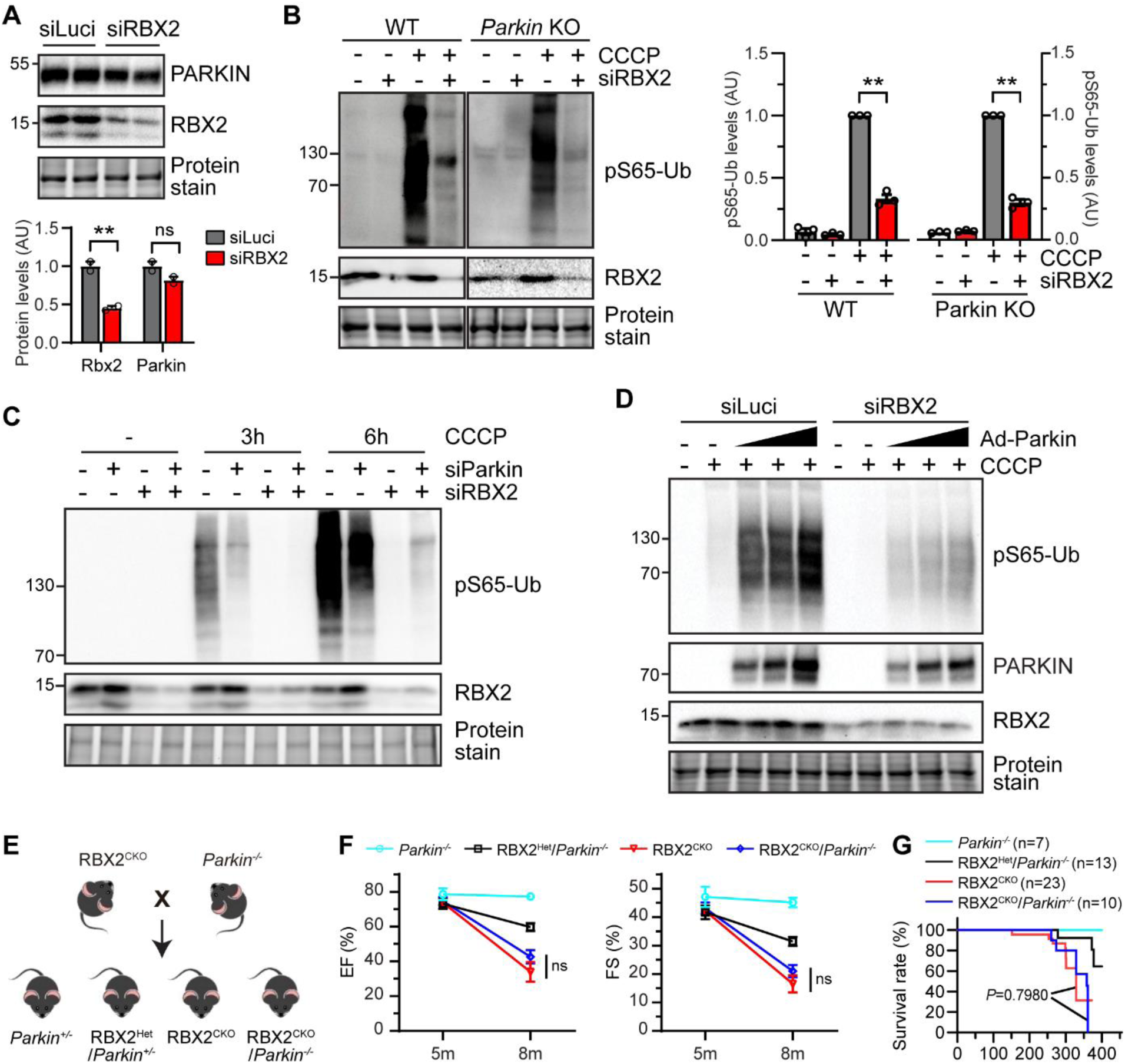
The role of Parkin in RBX2-regulated mitochondrial ubiquitination and cardiac homeostasis. **A**, Western blot of Parkin in NRVCs. Cells were infected with Ad-Parkin and transfected with indicated siRNAs. **B**, Western blots (left) and quantification (right) of pS65-Ub. Neonatal mouse ventricular CMs (NMVCs) were isolated from WT or *Parkin^-/-^* mouse hearts, transfected with siRNA, and treated with CCCP (10 µM) for 12 hours. **B**, Quantification of phosphorylated Ub. **C**, Western blots of cell lysates from NRVCs transfected with siRNAs and treated with CCCP (10 µM). **D**, Western blots of cell lysates from NRVCs infected with Ad-Parkin, transfected with siRNAs, and treated with CCCP (10 µM). **E**, Schematics of generation of RBX2 and Parkin double knockout (RBX2^CKO^/Parkin^-/-^) mice. **F**, Ejection fraction and fractional shortening at 5 (Parkin^-/-^: n=5, RBX2^Het^/Parkin^+/-^: n=8, RBX2^CKO^: n=12. RBX2^CKO^/Parkin^-/-^, n=8) and 8 (Parkin^-/-^: n=7, RBX2^Het^/Parkin^+/-^: n=7, RBX2^CKO^: n=11. RBX2^CKO^/Parkin^-/-^, n=8) months of age. **G**, Survival curves of indicated mice. **K**, A proposed model showing the role of RBX2-CRL5 in regulation of physiological mitophagy and cardiac homeostasis. Student *t* test was used in **A** and **B**. One-way ANOVA followed by post hoc Tukey test in **F**. Log-rank (Mantel-Cox) test in **G**. ** *P*<0.01. ns, not significant.

### RBX2 stabilizes PINK1

We next examined whether RBX2 regulates the phosphorylation of Ub at serine 65, which is known to be coordinated by PINK1 and phosphatases such as PPEF2, PGAM5 and PTEN-L 9-11. PINK1 is readily degraded by the proteasome in healthy mitochondria but quickly stabilized in depolarized mitochondria 12. Indeed, PINK1 expression was barely detected in cultured cardiomyocytes infected with adenovirus expressing PINK1 under basal condition (Figure 8A). Depletion of RBX2 significantly inhibited CCCP-induced PINK1 accumulation without affecting the expression of PPEF2, PGAM5 and PTEN (Figure 8A). Moreover, deletion of RBX2 in Hela cells via CRISPR/Cas9 also reduced CCCP-induced expression of endogenous PINK1 (Figure 8B). RBX2 deficiency did not alter the levels of PINK1 transcripts, and the reduction of PINK1 in RBX2-deficient cardiomyocytes was attenuated by the proteasome inhibitor bortezomib (Figure 8C and 8D), suggesting that RBX2 regulates the stability of PINK1. To determine whether RBX2 regulates PINK1 degradation, cardiomyocytes were treated with CCCP for 3 hours to promote PINK1 accumulation, and following removal of CCCP, PINK1 degradation was assessed by cycloheximide-based pulse chase assay. Under these conditions, results showed that loss of RBX2 shortened the half-life of PINK1 (Figure 8E). Moreover, in line with the reduced PINK1 expression in RBX2-deficient cardiomyocytes, RBX2 depletion increased the ubiquitinated forms of PINK1, which was blunted by proteasome inhibition (Figure 8F), indicating that RBX2 deficiency enhances the ubiquitination and degradation of PINK1. PINK1 mediates the phosphorylation of PARKIN, which promotes the translocation of PARKIN to mitochondria 12. Consistent with the reduced PINK1 expression, depletion of RBX2 abolished CCCP-induced PARKIN phosphorylation (Figure 8G), indicating diminished PINK1 activity. Consistent with the in vitro data, PINK1 expression was significantly reduced in RBX2iCKO hearts (Figure 6G). Together, these data suggest that RBX2 is required for PINK1 stabilization upon mitochondrial depolarization, which in turn enhances phosphorylation of Ub on mitochondria and thus mitochondrial ubiquitination.

**Figure 8.**
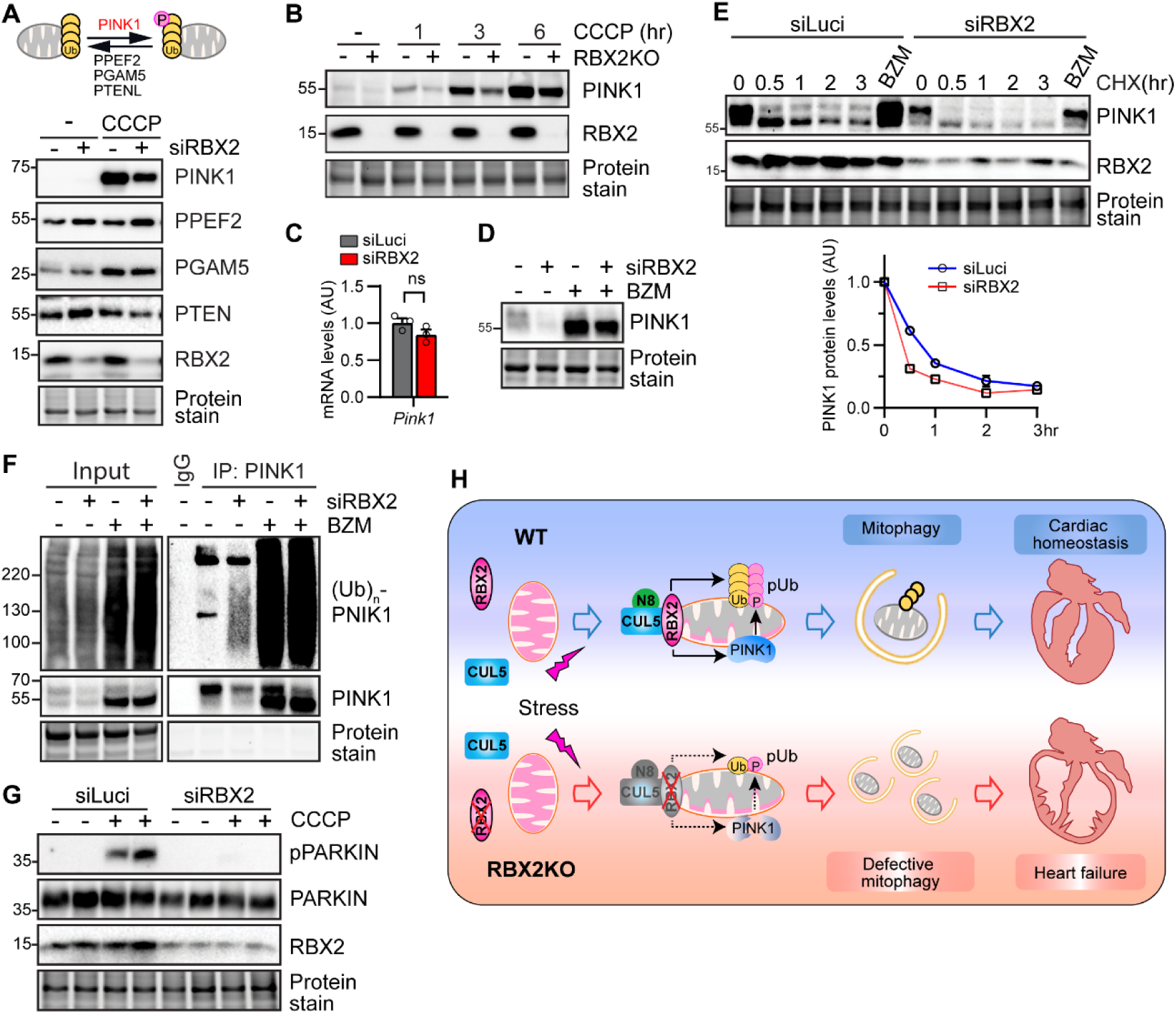
RBX2 stabilizes PINK1. **A**, Western blots of indicated proteins in NRVCs. Cells were infected with Ad-PINK1, transfected with indicated siRNAs and treated with or without CCCP (10 µM) for 12 hours. **B**, Western blots. *RBX2* was deleted in Hela cells via CRISPR/Cas9 using a single guided RNA against *RBX2* (gRBX2). WT and RBX2KO cells with treated with CCCP (10 µM) for indicated times before harvest. **C**, Analysis of *Pink1* transcript levels in NRVCs transfected with indicated siRNAs by qPCR. **D**, Western blots. NRVCs were infected with Ad-PINK1, transfected with siRNAs, and treated with Bortezomib (BZM, 100 nM) for 6 hours. **E**, Cycloheximide-based pulse chase assay. NRVCs were transfected with siRNAs, treated with CCCP (10 µM) for 3 hours, followed by removal of CCCP, and then chased for the indicated time in the presence of cycloheximide (CHX, 100 nM). **F**, Immunoprecipitation of PINK1, followed by Western blots. NRVCs were transfected with indicated siRNAs and treated with or without Bortezomib (BZM, 100 nM) for 6 hours. **G**, Western blots of cell lysates from NRVCs transfected with indicated siRNAs and treated with CCCP (10 µM) for 3 hours. **H**, A proposed model showing the role of RBX2-CRL5 in regulation of physiological mitophagy and cardiac homeostasis.

## DISCUSSION

In summary, our work identifies RBX2-CRL5 ubiquitin ligase as a novel regulator of mitophagy and cardiac homeostasis. Our evidence supports a model (Figure 8H) in which RBX2 translocates to dysfunctional mitochondria of cardiomyocytes, where it mediates the ubiquitination of mitochondrial outer membrane proteins and the clearance of damaged mitochondria under physiologic conditions in a PARKIN-independent manner. Meanwhile, RBX2 also enhances the stability of PINK1 to promote phosphorylation of Ub on depolarized mitochondria, thereby amplifying mitochondrial ubiquitination and mitophagy. Consequently, loss of RBX2 in mouse hearts impairs physiological mitochondrial turnover and provokes severe mitochondrial stress, leading to rapid development of dilated cardiomyopathy and heart failure. Our findings provide a new mechanism regulating mitochondrial quality control through targeting both mitochondrial ubiquitination and phosphorylation of ubiquitin on mitochondria. These findings may also explain why PARKIN is dispensable for physiological mitochondrial turnover in the heart.

### RBX2-CRL5 is a mitochondrial Ub ligase

RBX2-CRL5 participates in diverse cellular processes through targeting numerous substrates for ubiquitination^53^; however, its role in mitochondria has not been explored. Our studies identify RBX2-CRL5 as a mitochondrial Ub ligase. This is supported by the translocation of RBX2 and CUL5 to the mitochondrial outer mitochondrial membrane following mitochondrial depolarization, the necessity of RBX2 in mitochondrial ubiquitination and turnover, the global impact of RBX2 deficiency in mitochondrial proteome, and a crucial role of RBX2 in maintenance of mitochondrial integrity and function (Figure 1-2, Online Figure I-II). Retention of neddylated CUL5, an active form of CRL5, was noted in mitochondria (Figure 1C), further supporting its role in mitochondrial ubiquitination. In APEX2-MOM pulldown experiments, we observed more pronounced accumulation of RBX2 and CUL5 in mitochondrial fractions following CCCP treatment compared with crude mitochondrial fractions (Figure 1B-1C). These findings suggest that the interaction of RBX2-CRL5 with mitochondria is weak and transient, which may facilitate the rapid cycling of assembly and disassembly of CRL5 to recognize and ubiquitinate different MOM proteins by pairing with one of its 38 substrate-recognizing receptors. Since RBX2, CUL5 and its substrate receptors do not contain a mitochondrial targeting sequence, CRL5 may translocate to mitochondria through docking to unknown MOMs and/or other adaptor proteins. Interestingly, RBX2 was identified to be a redox sensitive protein, acting to scavenge reactive oxygen species at the cost of self-oligomerization via formation of intra- and inter-molecular disulfide bonds^54, 55^. Whether increased mitochondrial ROS might induce translocation of RBX2 to the mitochondria is unknown and worthy of future investigation. Furthermore, the molecular mechanisms whereby CRL5 controls mitochondrial ubiquitination including how CRL5 senses mitochondrial injury and interacts with mitochondria, which substrate receptors are involved in CRL5-mediated mitophagy, which mitochondrial proteins are the direct targets of CRL5, and whether its role in mitophagy is conserved in multiple cell types, remain to be elucidated. In line with the involvement of CRLs in mitophagy, two CUL3-associating adaptors, Keap1 and Klhl10, were reported to associate with mitochondria^56, 57^. The core component of CRL1-4, RBX1, was also reported to interact with mitochondria and may mediate mitochondrial ubiquitination in hepatocytes^58^. Moreover, FBXL4, a mitochondrial protein that forms the Skp1-Cullin 1-F-box (SCF)-FBXL4 ubiquitin ligase complex, prevents excessive mitophagy by mediating the degradation of mitophagy receptors BNIP and BNIP3L ^59^. Therefore, it is possible that other CRLs may also regulate mitophagy in cardiomyocytes.

### Essential role of RBX2 in regulation of physiological mitochondrial turnover and cardiac homeostasis

Evidence obtained through targeting key mitophagy regulators confirms an essential role of mitophagy in the heart under the conditions of stress ^60, 61^. In contrast, much less is known about how physiological mitophagy (mitochondrial clearance under steady-state conditions) is initiated and regulated in the heart. Our data indicate that, in normal hearts, only a small fraction of mitochondria undergo mitophagy at any given time, as evidenced by the colocalization of pS65-Ub or LAMP1 with mitochondria and mt-Keima reporter activity (Fig. 5A, 5G, 6D, and Online Figure VI). This suggests that mitophagy activity in normal hearts is relatively low, possibly as a safeguard to maintain overall mitochondrial homeostasis. However, accumulation of damaged mitochondria over time due to defective mitophagy can have catastrophic effects on mitochondrial and cardiac function. Deletions of general autophagy regulators, such as VPS34 and ATG5^62, 63^, or the key mitophagy regulator PINK1 ^64^ cause lethal cardiomyopathy and heart failure, supporting the importance of physiological mitophagy in cardiac homeostasis. In agreement with its pivotal role in mitophagy, loss of RBX2 in the postnatal and adult heart causes cardiac dysfunction and heart failure (Figure 3, Online Figure III-IV). The severe cardiac phenotype is attributable to mitochondria dysfunction arising from defective mitochondrial turnover, as evidenced by diminished mitochondrial ubiquitination, decreased mitophagy activity, accumulation of damaged mitochondria in the heart, compromised mitochondrial bioenergetics, and perturbations of cardiac metabolic pathways (Figure 5-6). Notably, RBX2 deficiency results in a rapid development of cardiomyopathy with much more pronounced cardiac dysfunction in adult hearts compared with developing hearts. The difference of disease progression rates in the two models could be due to a higher demand for energy production and thus mitochondrial homeostasis in adult hearts and/or the plasticity in adapting to mitochondrial dysfunction in young hearts, although we cannot completely exclude a confounding effect of tamoxifen-induced cardiotoxicity. Indeed, as seen in the cases of two ubiquitin ligases involved in mitophagy, TRAF2 and MITOL, and the key regulator ATG5, loss of these mitophagy regulators in adult hearts has a more profound impact on cardiac function, and typically leads to higher mortality, in adult mice than in developing mice^25, 63, 65, 66^. Notably, RBX1, the other RING-box family protein, was upregulated by 3-fold in CKO hearts but not in iCKO hearts (Figure 3B and Online Figure IVB), suggesting the possibility of compensation by RBX1 in the CKO heart, which awaits to be investigated in future experiments. Collectively, our data identify RBX2-CRL5 as an important regulator of physiological mitophagy and cardiac integrity. Nevertheless, considering that RBX2 controls the degradation of numerous protein substrates involved in various critical biological processes, including cell cycle, cytokine signal transduction, p53 signaling, mTOR signaling, and hypoxic response^53^, we cannot rule out the potential contribution of dysregulation in these pathways to the observed cardiac phenotypes in RBX2-deficient mice.

### Distinct actions of RBX2 and PARKIN in mitochondrial quality control

Despite the well-established role of PARKIN in mitophagy, it is increasing accepted that PARKIN is not the only initiator of mitochondrial ubiquitination and mitophagy ^67^. Given the low expression levels of PARKIN in the heart^12^ and the high abundance of mitochondria in cardiomyocytes, it is conceivable that multiple ubiquitin ligases cooperatively survey mitochondrial quality in the heart. Among the reported mitochondrial ubiquitin ligases that are involved in mitophagy, TRAF2 and MITOL have been showed to facilitate PARKIN recruitment to mitochondria and regulate mitophagy in the heart^26, 68^. We presented evidence demonstrating that RBX2 does not require PARKIN to mediate mitochondrial ubiquitination and possibly even acts downstream of PARKIN (Figure 7, Online Figure VIII). Moreover, loss of PARKIN does not impact cardiomyopathy induced by RBX2 deficiency, supporting a key role of RBX2 in physiological mitophagy in parallel to PARKIN. Several reports have also implicated PARKIN-independent roles for other cullin ubiquitin ligases in mitophagy. ARIH1, which belongs to Cullin 1-RING Ub ligase (CRL1) by forming a protein complex with Cullin 1 and RBX1, mediates mitophagy in PARKIN-deficient cancer cells ^20^. CUL9 ubiquitin ligase regulates mitochondrial quality control^69^, and loss of CUL9 and PARKIN together does not exaggerate mitochondria damage induced by CUL9 deficiency in neuronal cells ^70^. Additionally, the minimal requirement for mitochondrial ubiquitination, e.g., the number and amount of proteins that must be ubiquitinated, and the specific sites, necessary to trigger the autophagy machinery, remains to be determined. Given the well-documented role of PARKIN under stress conditions, it is possible that RBX2 and PARKIN may recognize different substrate pools (Online Figure VIIIE) to regulate mitophagy in cardiomyocytes under physiological and pathological conditions, respectively.

### RBX2 is a new regulator of PINK1 expression

An interesting finding of this study is that RBX2 is required for PINK1 stabilization following mitochondrial depolarization (Figure 8). PINK1 is imported into mitochondria and cleaved by the mitochondrial protease PARL^71^. Cleaved PINK1 is then degraded by the proteasome through the ubiquitin ligases UBR1/UBR2/UBR4^72^. It will be interesting in the future to explore whether RBX2 acts as a ubiquitin ligase to control the degradation of PARL or UBR1/2/4, thereby increasing PINK1 stability in damaged mitochondria. Loss of PINK1 promotes mitochondrial dysfunction in the heart eventually leading to cardiac dysfunction and heart failure^64^. The dual role of RBX2 in regulation of mitochondrial ubiquitination and PINK1 expression explains the severe mitochondrial and cardiac phenotypes observed in adult RBX2-deficient hearts.

## METHODS and MATERIALS

### Animals

Cardiomyocyte-specific RBX2 knockout (RBX2^CKO^) mice were generated by crossing a *Rbx2^Flox^* allele with Exon 1 flanked with loxP sites^37^ with *αMHC^Cre/+^* mice (the Jackson Laboratory, strain #011038). Inducible cardiomyocyte-specific RBX2 knockout (RBX2^iCKO^) mice were generated by crossing *Rbx2^F/+^* mice with αMHC-MerCreMer mice (MCM, the Jackson Laboratory, strain # 005657). To induce RBX2 knockout in adulthood, 12 to 16-week-old mice were treated with tamoxifen (Sigma, Cat# T5648) by intraperitoneal injection at a dosage of 50 mg/kg/day for 5 days or 20 mg/kg/day for 10 days (5 consecutive days with a 2-day interval in between). Tamoxifen was first dissolved in 5% ethanol and further diluted in sunflower oil (Sigma, Cat# S5007). *Parkin* knockout mice (B6.129S4-Prkn*^tm1Shn^*/J) were obtained from The Jackson Laboratory (strain #006582). All mice were maintained on a C57BL/6J background. Primers used for genotyping are listed in Online Table III. All animal experiments were approved by the Augusta University Institutional Animal Care and Use Committee.

### APEX2-MOM labeling and biotinylated proteins enrichment

This was accomplished as previously reported^39^ with minor modification. NRVCs were infected with adenovirus expressing APEX2-MOM (VectorBuilder). After 48 hours, cells were treated with or without CCCP (10 µM) for different times and subsequently cultured in fresh growth medium containing 500 µM biotin-phenol (AdipoGen Life Sciences) at 37°C under 5% CO_2_ for 30 minutes, followed by the addition of H_2_O_2_ to a final concentration of 1 mM H_2_O_2_ for 1 minute at room temperature on a rotator with gentle agitation. The reaction was then quenched by washing three times with Dulbecco’s PBS (DPBS) containing 5 mM Trolox, 10 mM sodium azide, and 10 mM sodium ascorbate. Cells were then lysed in RIPA lysis buffer (50mM Tris, 150 mM NaCl, 0.1% SDS, 0.5% sodium deoxycholate, 1% Triton X-100, 5 mM EDTA, protease cocktail, 1 mM PMSF, 10 mM sodium azide, 10 mM sodium ascorbate, and 5mM Trolox) for 10 min at 4°C, followed by sonication. Cell lysates were clarified by centrifugation at 12,000 g for 10 min at 4°C. The pellets were resolubilized in 6 M urea and 1% SDS and then combined with the soluble fraction. Protein lysates (500 μg) from each biological sample were incubated with 100 μL of NeutrAvidin Agarose Resins (Thermo Fisher, Cat# 29200) with rotation for 1 hour at room temperature. The beads were subsequently washed twice with 1 mL RIPA lysis buffer, once with 1 mL of 1 M KCl, once with 1 mL of 0.1 M Na_2_CO_3_, once with 1 mL of 2 M urea in 10 mM Tris-HCl pH 8.0, and twice with 1 mL RIPA lysis buffer. Biotinylated proteins were eluted by boiling the beads in 30 µl of 2X SDS-PAGE sampling buffer supplemented with 50 mM DTT and 50 μM biotin for 10 minutes. The beads were eluted twice, and the eluents were combined for subsequent Western blotting.

### Assessment of mitophagy activity

Mitophagy in live cells was monitored using the Mitophagy detection kit (Dojindo Molecular Technologies) according to the manufacturer’s instructions. Briefly, cells were incubated with 100 nM Mtphagy Dye (Excitation at 547 nm) working solution for 30 minutes, then treated with 10 μM CCCP to induce mitophagy for 3 hours, and incubated with 1 µM LysoTracker (Excitation at 405 nm) Dye (Life Technologies) for 30 minutes at 37°C. The level of mitophagy activity was defined by the number of Mtphagy-positive puncta per cell.

To evaluate mitophagy activity with the mt-Keima reporter *in vivo*, neonates at age P2-P3 were injected subcutaneously with AAV-mt-Keima (Charles River, 1X10^11^ GC/pup), in which the expression of mt-Keima was driven by the CAG promoter. At 12 weeks of age, intact mouse hearts were excised and rinsed with PBS. The mt-Keima signals in epicardial myocytes were detected at 488 nm and 568 nm *in situ* with confocal microscope (STELLARIS 8, Leica Microsystems). The heatmaps representing 568/488 ratio were analyzed and generated using MATLAB software. Specifically, the intensity of each pixel within the 568 and 488 channel images was first exported to data matrixes using “imread” function. Next, appropriate thresholds were identified for each channel to eliminate background signals. After the 568/488 ratio of each pixel was calculated, the ratio matrix was remapped into a heatmap using “colormap” function in ‘jet’ mode, and color scales using “colorbar” function. Images from different views (typically 20-40 views per heart, 3 hearts per group) were included in the quantification. For each view, the ratio of intensity of 568/488 channels was quantified in ImageJ (NIH) and plotted with Graphpad Prism.

To assess cardiac mitophagy flux by immunoblotting, mice were intraperitoneally injected with bafilomycin A1 (BFA; 3μmol/kg, Sigma). BFA was dissolved in 50% DMSO. Hearts were collected 3 hours after the injection for mitochondrial and cytosolic fractionation and subsequent assessment of LC3-II protein levels by Western blot.

### Peptide preparation, TMT labelling, and mass spectrometry

This was performed by following a previously reported protocol ^73^. Cells were lysed with ice cold urea lysis buffer [8M urea,75mM NaCl, 50mM pH 8.0 Tris-HCl, 1mM EDTA, and 50 μM PR-619 (LifeSensors, SI9619) with proteinase inhibitor cocktails (Roche, Cat# 04693159001)]. Protein lysates were diluted to 2 M urea with 50 mM HEPES (pH 8.2), The concentration of protein lysates was determined. Equal amount (50 µg) of proteins from each biological sample were reduced by 10 mM TCEP (Thermo Fisher Scientific, Ca# 20490) and alkylated by 20 mM iodoacetamide (Thermo Fisher Scientific, Ca# A39271) in the dark for 45 min at 45 °C. The protein lysates were then digested with trypsin (Thermo Fisher Scientific, Ca# 90059) at a 1:50 enzyme/protein ratio overnight at 37°C with gentle end-over-end rotation. The concentration of the peptides was determined using Quantitative Colorimetric Peptide Assay (Thermo Fisher Scientific, Cat#23275). Tandem mass tag (TMT)-based labeling of peptides was performed using the TMT10plex™ Isobaric Label Reagent Set (Thermo Fisher Scientific, Cat#90110) following the manufacturer’s instructions. Briefly, 2 μL of a TMT10 reagent (0.2 mg/11 μL) was incubated with 4.5 µg peptides from each biological replicate at room temperature for 1 hour with shaking (700 rpm). The reaction was quenched with 2 μL of 5% hydroxylamine for 5 minutes. TMT-labeled samples were combined and desalted using 100 µl Pierce® C18 Tips (Thermo Fisher Scientific, Cat#87784) and dried in an Vacufuge concentrator (Eppendorf 5301). The resultant peptides (a total of 45 ug from 10 samples) were sent to the Center for Mass Spectrometry and Proteomics (CMSP) at the University of Minnesota for mass spectrometry analysis.

For mass spectrometry analysis, the peptide was reconstituted in 2% acetonitrile/0.1% formic acid (FA) and analyzed by an Orbitrap Eclipse Tribrid mass spectrometer (Thermo Fisher Scientific) coupled online to an UltiMate 3000 RSLCnano System (Thermo Fisher Scientific). Briefly, 5 μg of sample was loaded onto a microcapillary column (360 μm OD×75μm ID) containing an integrated electrospray emitter tip (10 μm) (New Objective) packed with 30 cm of ReproSil-Pur C18-AQ 1.9 μm beads (Dr. Maisch GmbH). The nanoflow column was heated to 55 °C using a column heater (Phoenix S&T). Samples were analyzed using a 199 min LC-MS method. Mobile phase flow rate was 200 nL/min. Solvent A was comprised of 0.1% FA. Solvent B was comprised of 0.1% FA in acetonitrile. The LC-MS/MS method used the following gradient profile: (min: %B) 0:2; 1:6; 122:35; 130:60; 133:90; 143:90; 144:50; 154:50 (the last two steps at 500 nl/min flow rate). The mass spectrometer was operated such that MS1 spectra were measured with a resolution of 60,000, an AGC target of 4 × 105, and a mass range from 350 to 1800 m/z. A top speed approach (cycle time 2 s) was used to trigger MS/MS at a resolution of 50,000, an AGC target of 1 × 105, an isolation window of 0.7 m/z, a maximum ion time of 150 ms, and a normalized collision energy of 38. The dynamic exclusion time was set to 20 s, the peptide match was set to peptide mode, and charge state screening was enabled to reject precursor charge states that were unassigned, 1, or >6.

For data analysis, the false discovery rate (FDR) was set at 0.01 for the identification of peptides and proteins. All the proteins were identified by two or more unique peptides. Reporter ion isotopic distribution correction was pursued in order to correct the factors for natural carbon isotopes and incomplete isotope incorporation. The reporter ion isotopic distributions obtained from the product data sheet (WA306750) of the TMT reagent were used as isotope correction factors in TMTpro 10plex method template using in-house Proteome Discoverer software (version 2.5.0.400). Differentially expressed proteins were defined by a threshold of P<0.05 and a fold change of 1.3. Pathway enrichment was performed using Metascape^74^.

### RNA-seq analysis

Mouse ventricles were minced and treated with RNAlater (Thermo Fisher Scientific). Total RNA was isolated using TRIzol reagent (Thermo Fisher Scientific) according to the manufacturer’s instructions. The isolated RNA was subjected to RNA-sequencing analysis performed by Novogene via Next Generation Sequencing. Differentially expressed genes were defined by *P*adj<0.05 and a fold change larger than 1.5. Pathway enrichment analysis was carried out using Metascape^74^ with a nominal threshold of *P*<0.05.

## Supporting information

Supplemental manuscript

## ACKNOWLEDGEMENTS

This study was in part supported by the US National Institutes of Health grants (R01HL124248 and R01HL165205 to H.S. and R01HL146807A1 to J.L.) and the American Heart Association grant (959479 to H.S.).

## CONTRIBUTIONS

W.W. and H.S. conceptualized the project and designed the experiments. W.W., E.L., J.Zou., Q.C., M.I., J.A. and Y.W. performed the experiments; W.W., E.L., J.Zou., Q.C., J.A., Y.W., J.Li and H.S. performed the analysis of the data; N.L.W., J.Liu, J.Zhou, Q.Liang, J.Li, Y.S. and H.S. provided resources; W.W. and H.S. wrote the original draft and the final version of the manuscript; N.L.W., D.J.F., J.Li and Y.S. wrote, reviewed and edited the manuscript; H.S. supervised the project. All authors read and approved the final paper.

## Non-standard abbreviations and acronyms

RBX2: RING-Box Protein 2
SAG: Sensitive to Apoptosis Gene
Ub: Ubiquitin
pS65-Ub: phosphorylated Ub at serine 65
MAVS: mitochondrial antiviral-signaling protein
AAV: adeno-associated virus
AV: adenovirus
siRNA: Small interfering RNA
GFP: green fluorescent protein
CUL: cullin
RING: Really Interesting New Gene
CRLs: cullin-RING ligases
CSN: COP9 signalosome
APEX2: ascorbate peroxidase 2
mito: mitochondrial
cyto: cytosolic
MOM: mitochondrial outer membrane
CCCP: Carbonyl Cyanide Chlorophenylhydrazone
OMP25: Outer membrane protein 25
PK: proteinase K
HA: hemagglutinin
TMRM: Tetramethylrhodamine methyl ester perchlorate
αMHC: α- myosin heavy chain
CKO: cardiomyocyte-specific knockout
TAM: tamoxifen
TMT: tandem mass tag
KD: knockdown
CTL: control
MCM: MerCreMer
iCKO: inducible cardiomyocyte-specific knockout
BFA: bafilomycin A1
PCA: principle component analysis
MS: Mass spectrometry
DEPs: differentially expressed proteins
FC: fold change
FDR: False Discovery Rate
KEGG: Kyoto encyclopedia of genes and genomes
ER: endoplasmic reticulum
DKO: double knockout
CM: cardiomyocyte
cTnT: cardiac troponin T
NRVCs: neonatal rat ventricular cardiomyocytes
NRVMs: neonatal mouse ventricular cardiomyocytes
NMVFs: neonatal mouse ventricular fibroblasts
HF: heart failure
KO: knockout
MF: Molecular Functions
CC: Cellular Components
BP: Biological Process
TUNEL: Terminal deoxynucleotidyl transferase dUTP nick end labeling
SCF: Skp1-Cullin 1-F-box.

