## Supplemental manuscript for "The Ubiquitin Ligase RBX2/SAG Regulates Mitochondrial Ubiquitination and Mitophagy"

**Short Title**: RBX2 regulates mitophagy

**This supplementary information includes:**

Online Figures and Figure Legends, Supplemental Material and Methods, Supplemental References

**Online Figure I**

Association of CRLs with mitochondria.

**Online Figure II:**

Loss of RBX2 inhibits mitochondrial ubiquitination and exacerbates stress-induced cardiomyocyte cell death.

**Online Figure III**:

Deletion of Rbx2 in the adult heart by low dose of tamoxifen causes cardiac dysfunction.

**Online Figure IV**:

Cardiac-specific deletion of *Rbx2* leads to cardiomyopathy

**Online Figure V**:

RBX2 regulates the expression of mitochondrial proteins.

**Online Figure VI**:

Assessment of mitophagy in RBX2CKO hearts with mt-Keima reporter.

**Online Figure VII**:

Transcriptomic alterations in RBX2$^{iCKO}$ hearts.

**Online Figure VIII**:

Parkin is not required for the action of RBX2 in mitophagy and in maintenance of cardiac function.

**Online Table 1**. Differentially expressed genes identified between control and RBX2$^{iCKO}$ hearts.

**Online Table 2**. Differentially expressed proteins identified between siLuci and siRBX2-transfected cardiomyocytes.

**Online Table 3**. Oligonucleotides, antibodies and compounds used in this study.

**Online Table 4.** Echocardiography of RBX2 iCKO mice receiving tamoxifen injection (50 mg/kg).

**Online Table 5.** Echocardiography of RBX2 iCKO mice receiving tamoxifen injection (20 mg/kg).

**Online Table 6.** Echocardiography of RBX2 CKO mice.

**Online Table 7.** Echocardiography of Parkin and RBX2 double knockout mice.

#### Online Figures and Figure Legends

#### Online Figure I

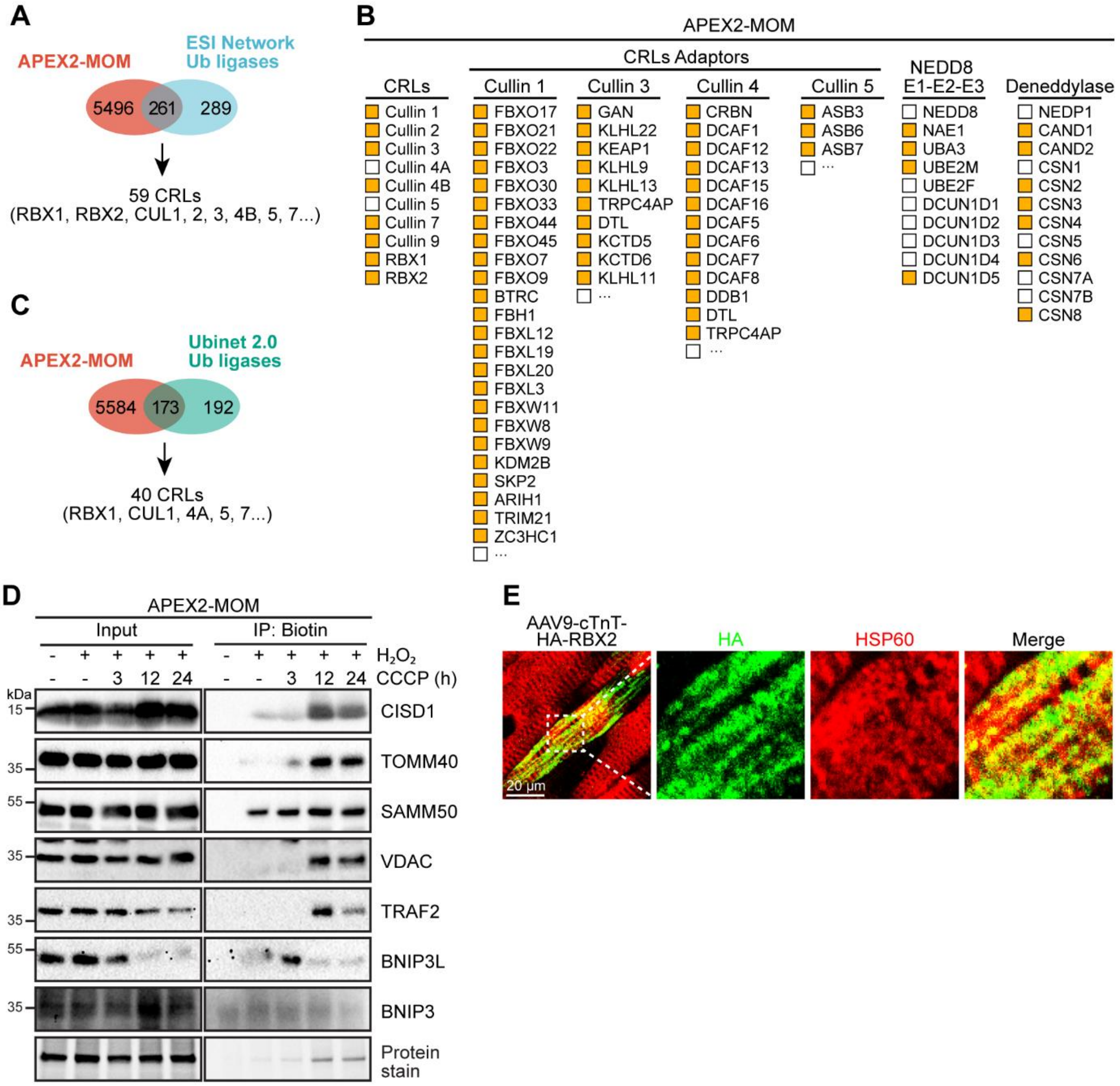


**Online Figure I**. **Association of CRLs with mitochondria. A**, Venn diagram showing the overlap of mitochondria-interacting proteome with the Ub ligases annotated in ESI Network. **B**, A list of CRLs components and neddylation enzymes identified by APEX2-MOM (yellow). **C**, Venn diagram showing the overlap of mitochondria-interacting proteome with the Ub ligases annotated in Ubinet 2.0. **D**, Representative Western blot of biotinylated proteins in neonatal rat ventricular cardiomyocytes (NRVCs) with adenoviral (Ad) expression of APEX2-MOM. NRVCs were treated with CCCP (10 µM) before $H_2O_2$ activation. **E**, Representative confocal images of myocardium sections from mice transduced with AAV9 expressing HA-RBX2 under the control of TnT promoter. HA (green) and HSP90 (red) were used to label RBX2 and mitochondria, respectively.

#### Online Figure II

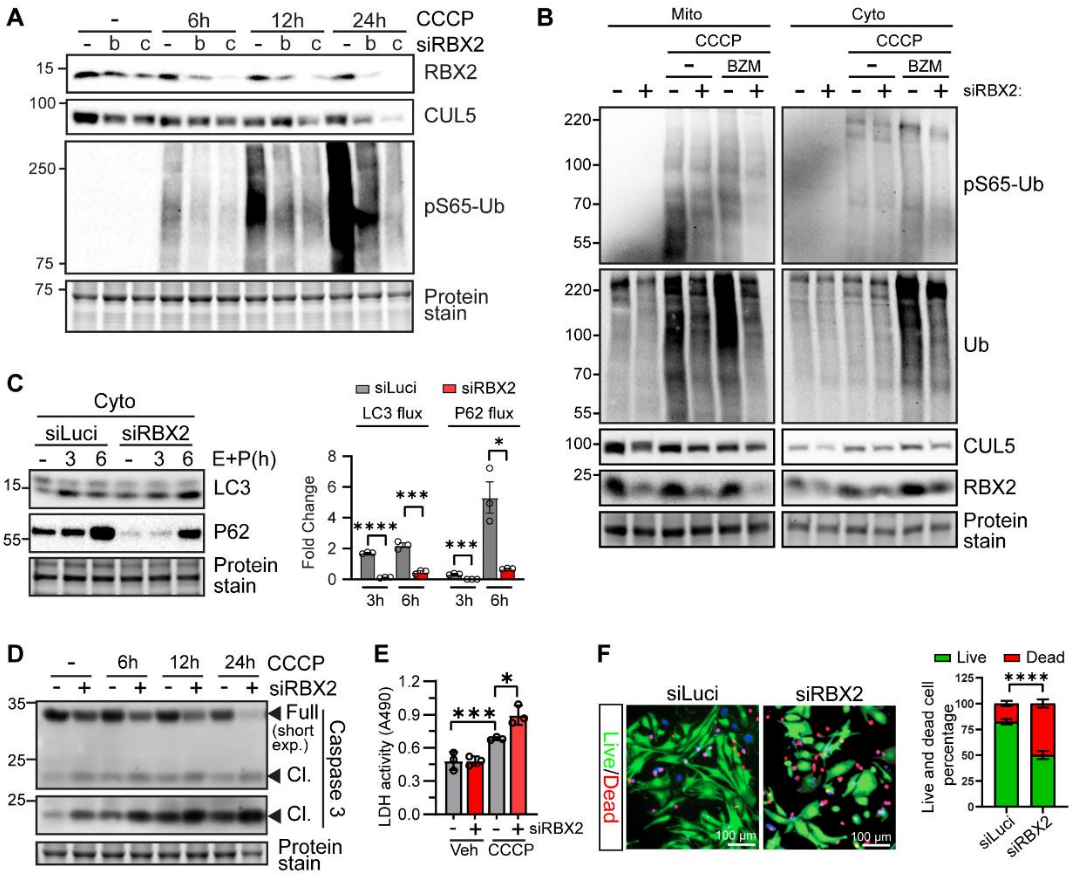

**Online Figure II**: **Loss of RBX2 inhibits mitochondrial ubiquitination and exacerbates stress-induced cardiomyocyte cell death**. **A**, Western blot of total cell lysates from NRVCs transfected with two different siRBX2s (b and c). CCCP (10 µM) were added for the indicated times. **B**. Western blots of mitochondrial (mito) and cytosolic (cyto) fractions. NRVCs were transfected with indicated siRNAs, treated with CCCP (10 µM) in the presence or absence of Bortezomib (100 nM) for 6 hours. **C**, Western blot of cytosolic extracts showing the impact of RBX2 deficiency on autophagy flux. NRVCs were treated as described in **Figure 2E**. **D**, Western blot of Caspase 3 in NRVCs. Note the reduction of full-length Caspase 3 and the corresponding increased cleaved (Cl.) form. **E**, Lactate dehydrogenase (LDH) activity in NRVCs at 3 hours after CCCP (10 µM) treatment. **F**, Live (green)/dead (red) cell staining of NVRCs exposed to CCCP (10 µM) for 6 hours. Representative fluorescent images (left) and the quantification (right) are shown. Student *t* test was used. * *P* <0.05, ** *P*<0.01, *** *P*<0.001, **** *P* <0.0001.

**Online Figure III**

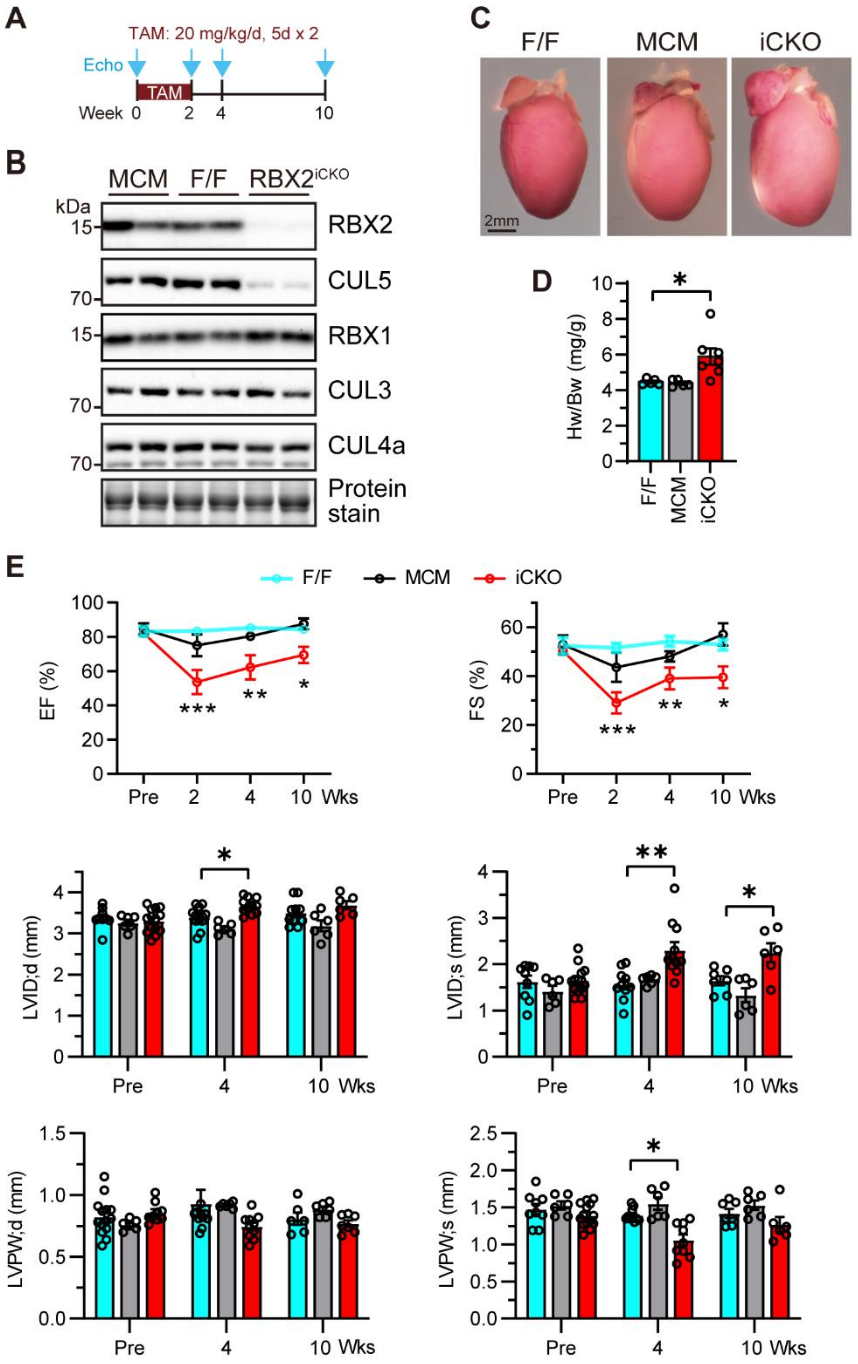


**Online Figure III. Deletion of *Rbx2* in adult heart by low dose of tamoxifen causes cardiac dysfunction. A**, Schematic experimental strategy for inducible RBX2 knockout by tamoxifen injection intraperitoneally at 20 mg/kg per day for 10 days wither a two-day interval after the first 5 injections. Heart function was measured by echocardiography before and after the tamoxifen injection. **B**, Representative immunoblot of RBX2 and other Cullin proteins at 15 days after tamoxifen treatment. **C**, Gross morphology of hearts at 10 weeks after tamoxifen injection. **D**, Heart weight to body weight ratio (Hw/Bw). F/F: n=5, MCM: n=5, iCKO: n=7. **E**, Quantification of echocardiographic parameters before and 2, 4, and 10 weeks post-tamoxifen injection. F/F: n=10, MCM: n=6, iCKO: n=9. One-way ANOVA followed by post hoc Tukey test was used in **D** and **E**. * $P$ <0.05, ** $P$<0.01, *** $P$<0.001.

#### Online Figure IV

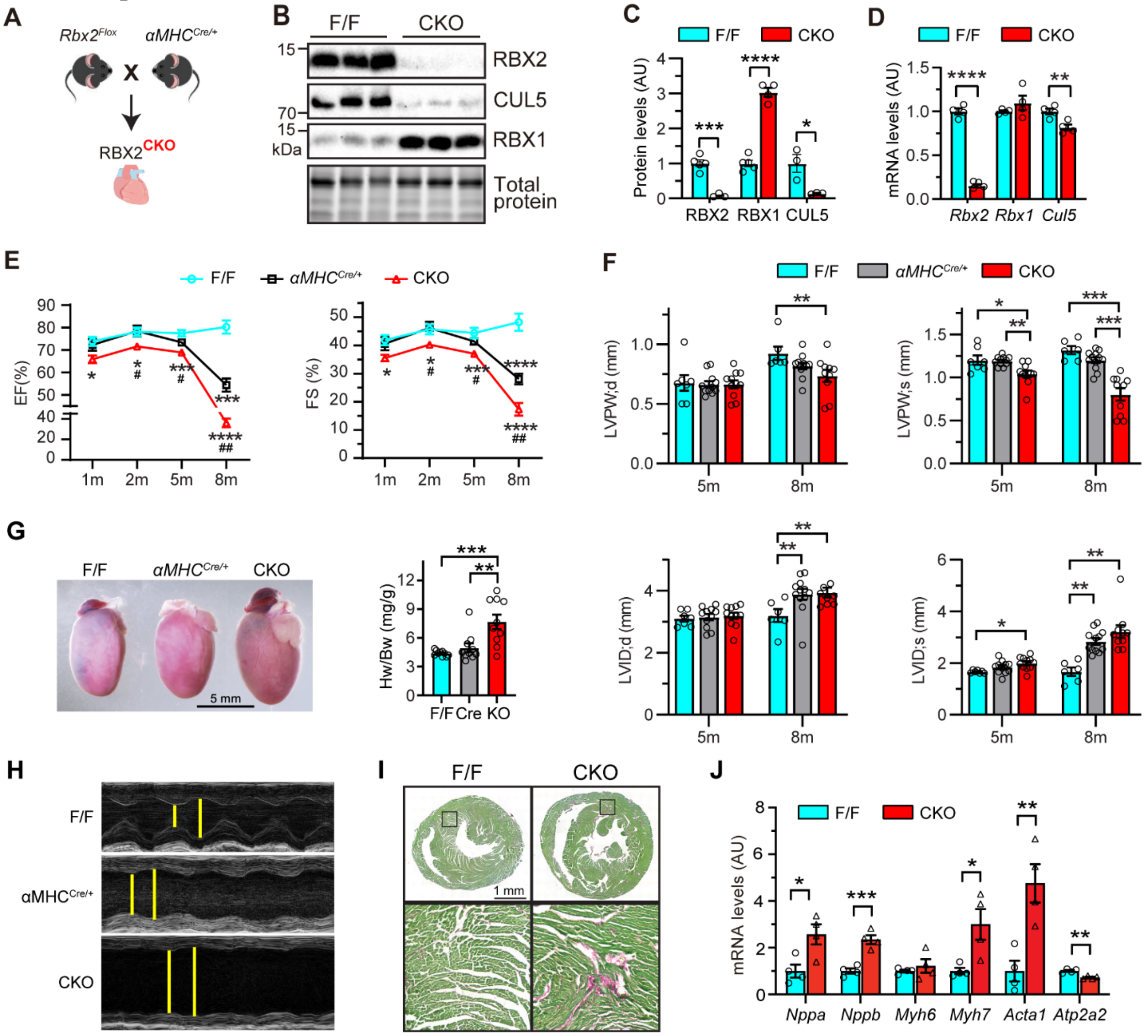


**Online Figure IV. Cardiac-specific deletion of *Rbx2* leads to cardiomyopathy and heart failure. A**, Schematic of generation of *αMHC*$^{Cre}$-mediated, cardiac-specific RBX2 knockout (CKO) mice. **B-C**, Western blot and quantification of indicated proteins in 2-month (m)-old hearts. F/F: n=4, CKO: n=3. **D**, Relative mRNA levels of the indicated genes in 2-month-old mouse hearts. F/F: n=3, CKO: n=3. **E**, Temporal changes in ejection fraction (EF) and fractional shortening (FS). F, Changes in Left ventricular posterior wall, diastolic (LVPW,d), Left ventricular posterior wall, systolic (LVPW,s), Left ventricular internal diameter, diastolic (LVID,d) and Left ventricular internal diameter, systolic (LVID,s). **G**, Gross heart size (left) and the quantification of heart weight (Hw) to body weight (Bw) ratio at 8 months of age. F/F: n=10, αMHC$^{Cre}$: n=10, CKO: n=10. **F**, Quantification of echocardiographic parameters at 5 (F/F: n=7, αMHC$^{Cre}$: n=11, CKO: n=11) and 8 (F/F: n=6, αMHC$^{Cre}$: n=11, CKO: n=10) months of age. **H**, Representative B-mode images at 8 months of age. Left ventricular internal diameters at systole and diastole are marked. **I**, Representative images of myocardial sections stained with Fast Green and Direct red 80 (PSR). **J**, qPCR analysis of indicated genes in 8-month-old hearts. F/F: n=4, CKO: n=4. Student *t* test was used in **B**, **C**, and **J**. One-way ANOVA followed by post hoc Tukey test in **D**, **E** and **F**. * $P<0.05$, ** $P<0.01$, *** $P<0.001$, **** $P<0.0001$.

**Online Figure V.**

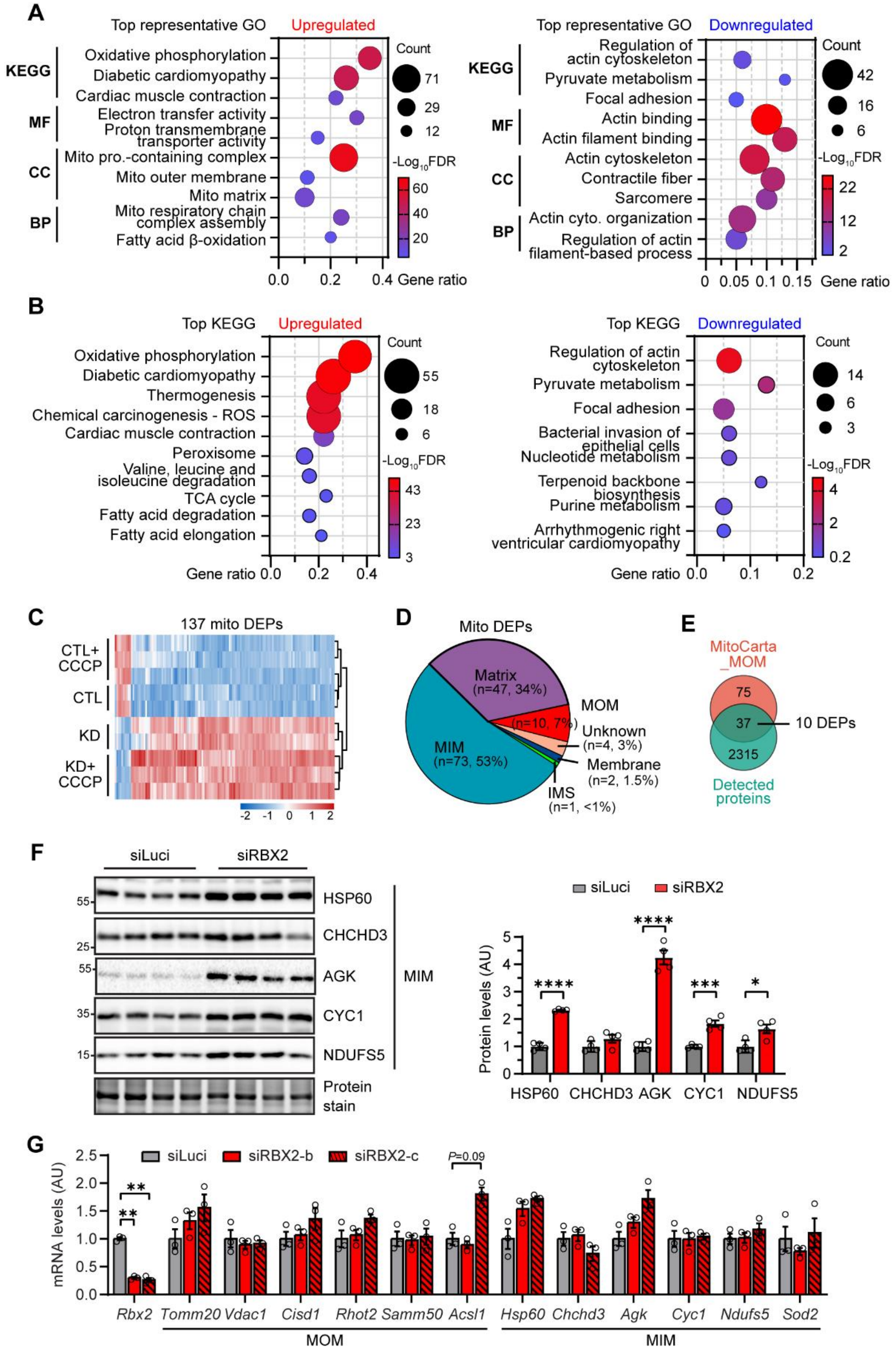


**Online Figure V. RBX2 regulates the expression of mitochondrial proteins. A**, Gene ontology analysis of upregulated (left) and downregulated (right) DEPs. **B**, Top

KEGG pathways enriched in upregulated (left) and downregulated (right) proteins identified in CCCP-treated, RBX2-deficient cardiomyocytes. **C**, Heatmap showing the expression of 137 mitochondrial proteins differentially expressed in RBX2-deficient cardiomyocytes among 4 groups. **D**, Venn diagram showing the distribution of 137 mitochondrial DEPs in different mitochondrial compartments. **E**, Venn diagram showing the overlap of MOMs annotated in MitoCarta 3.0 with proteins quantitatively detected in this study. Among 37 MOMs detected in this study, 10 were dysregulated in RBX2-deficient cardiomyocytes. **F**, Western blot (left) and quantification (right) of mitochondrial inner membrane proteins showing their accumulation in RBX2-deficient cardiomyocytes. n=4 for each group. **G**, QPCR analysis of indicated mitochondrial genes. n=3 for each group. Multiple *t* test was used. * $P$ <0.05, ** $P$<0.01, *** $P$<0.001, and **** $P$<0.0001.

#### Online Figure VI

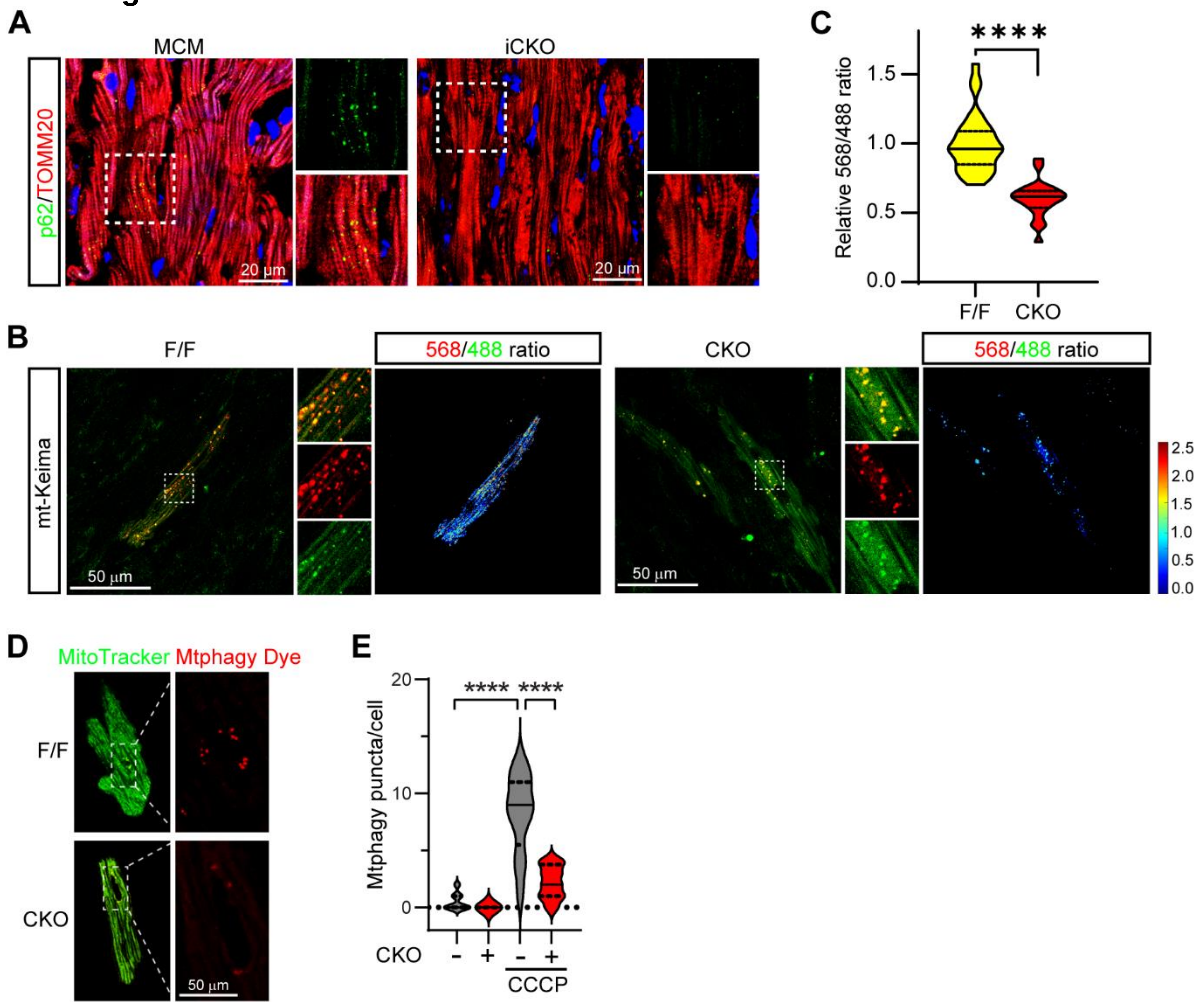


**Online Figure VI. Mitophagic activity in RBX2-deficient hearts**. **A**, Representative confocal images of MCM and RBX2$^{iCKO}$ myocardium sections immunostained with p62 (green), TOMM20 (red, mitochondrial marker) and DAPI. Scale bars, 20 µm. **B**, Representative confocal images of mt-Keima at 488 nm and 568 nm, respectively, in epicardial cardiomyocytes from RBX2$^{CKO}$ and littermate control (RBX2$^{F/F}$) mouse hearts and the derived heatmaps. Neonatal mice were infected with AAV-mt-Keima ($1X10^{11}$ GC/pup). At 2 months of age, hearts were excised and scanned for the mt-Keima signals in epicardial cardiomyocytes *in situ* with confocal microscope. Scale bars, 20 µm. **C**, Quantification of relative 568/488 ratio per cell. 20-40 views per heart, 2 hearts per group were quantified. **D,** Representative confocal images of live adult cardiomyocytes stained with Mtphagy dye (red) and MitoTracker (green) showing mitophagic vesicles. Adult cardiomyocytes were isolated from 2-month-old F/F and CKO mice. **E**, Quantification of mitophagic puncta per cell. Over 20 adult cardiomyocytes per group were quantified. Nested *t* test was used in **C**, and One-way ANOVA in **E.** **** $P$ <0.0001.

### Online Figure VII

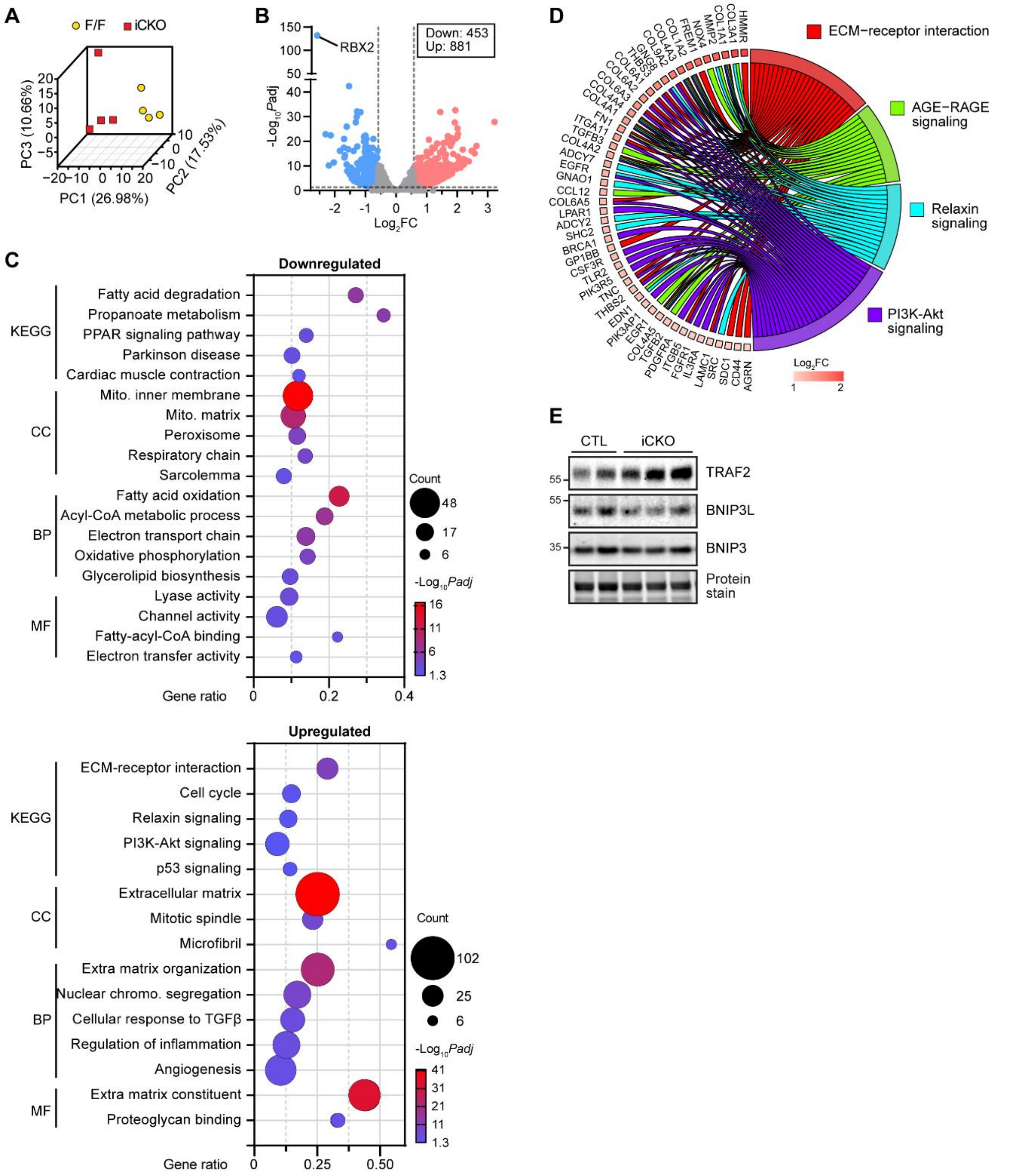


**Online Figure VII. Transcriptomic alterations in RBX2$^{iCKO}$ hearts. A**, Principal component analysis. F/F, n=4, iCKO: n=4. **B**, Volcano plot showing differentially expressed genes detected in iCKO hearts. **C**, Gene ontology analysis revealing top KEGG pathway, biological processes (BP), cellular components (CC) and molecular function (MF) enriched in downregulated and upregulated genes. **D**, Chord plot showing upregulated genes enriched in the indicated KEGG pathways. **E**, Western blot of indicated proteins in mouse hearts at 12 days after tamoxifen injection.

**Online Figure VIII**

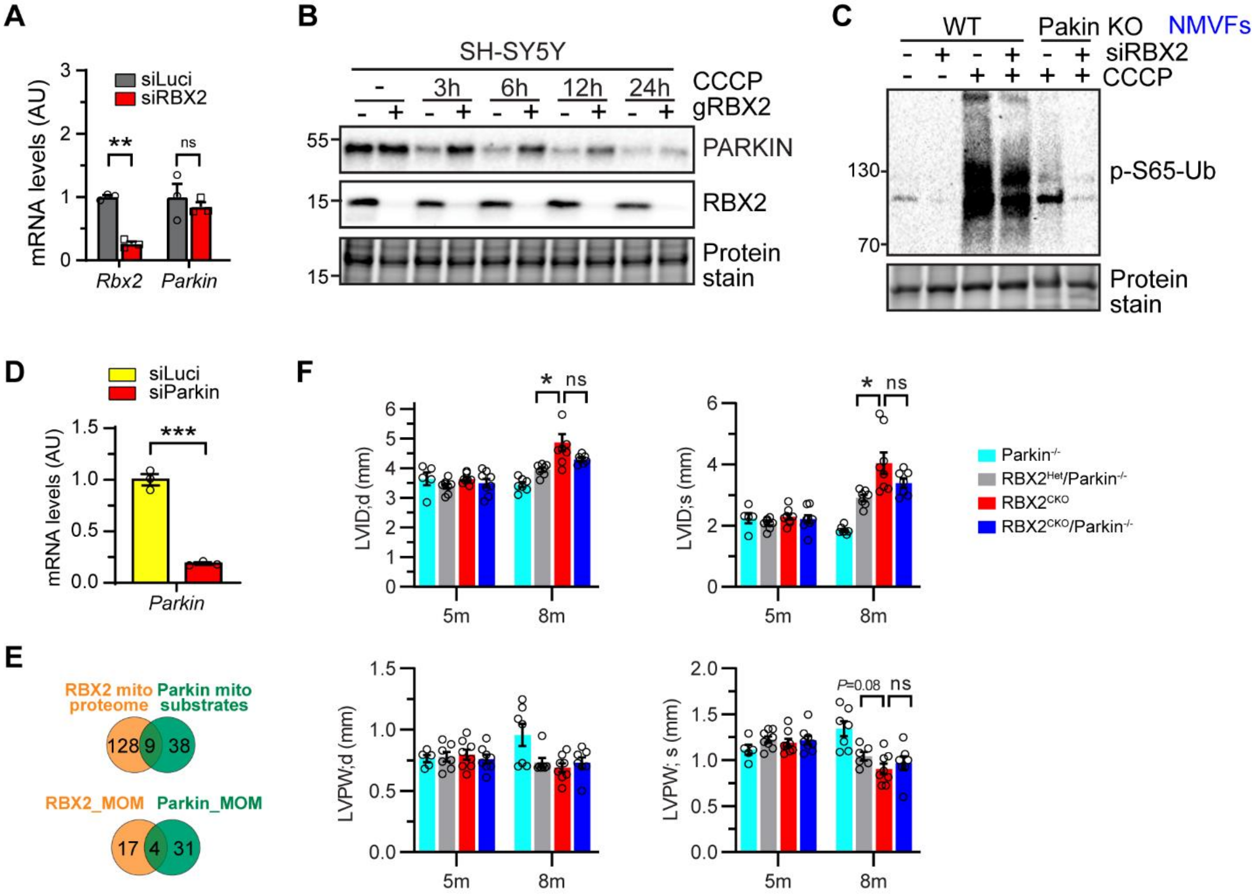


**Online Figure VIII**. **Parkin is not required for the action of RBX2 in mitophagy and in maintenance of cardiac function**. A, QPCR analysis of *Parkin* mRNA levels in RBX2-deficient cardiomyocytes. n=3 per group. B, Western blot showing the effect of RBX2 deficiency on the levels of PARKIN proteins in SH-SY5Y cells. The gene encoding RBX2 was deleted in SH-SY5Y cells via CRISPR/Cas9 using guided RNA against *RBX2* (gRBX2). The cells were treated with CCCP (10 µM) before analysis. **C**, Western blot showing the impact of RBX2 depletion via siRNAs on the levels of pS65-Ub in cardiac fibroblasts isolated from WT and Parkin$^{-/-}$ hearts. **D**, Validation of *Parkin* expression in NRVCs transfected with siRNA against *Parkin* by qPCR. **E**, Venn diagrams showing the overlap of RBX2- and Parkin-regulated mitochondrial substrates (top) and MOMs (bottom). **E**, Quantification of echocardiographic parameters at 5 months (Parkin$^{-/-}$: n=5, RBX2$^{Het}$/Parkin$^{+/-}$: n=8, RBX2$^{CKO}$: n=12. RBX2$^{CKO}$/Parkin$^{-/-}$, n=8) and 8 (Parkin$^{-/-}$: n=7, RBX2$^{Het}$/Parkin$^{+/-}$: n=7, RBX2$^{CKO}$: n=11. RBX2$^{CKO}$/Parkin$^{-/-}$, n=8) months of age. Student *t* test was used in **A**. One-way ANOVA followed by post hoc Tukey test in **E**. * *P*<0.05. ns, not significant.

**Supplemental Material and Methods**, **Supplemental References**

#### Culture of neonatal cardiomyocytes and cell lines

Neonatal rat or mouse ventricular cardiomyocytes (NRVCs or NMVCs) were isolated using Neonatal Cardiomyocyte Isolation System (Worthington Biochemicals, Cat# LK003245) following the manufacturer's protocol. Briefly, hearts from neonatal rats or mice were excised with the removal of the atrium, minced into ~1 mm diameter, and digested in 0.05% Trypsin in 4°C overnight on a slow-speed rocker. The next day, the heart tissues were washed 4 times and subjected to collagenase digestion at 37°C for 40 min. Cardiomyocytes were next separated by pipet flushing and through 70 µm filter, pre-plated for 2 hour to exclude cardiac fibroblasts, and finally plated in 60-mm dishes in DMEM containing 10% FBS, 1% P/S and 1x BrdU for two days in 37°C incubator with 5% CO2 prior to use for subsequent experiments.

For genetic knockdown and overexpression experiments, cardiomyocytes were transfected with siRNA of interest using Lipofectamine RNAiMAX (Thermo Fisher Scientific) following the manufacturer's protocol or infected with adenovirus encoding gene of interest as we previously reported 2. The small interference RNA (siRNA) used in this study were designed and synthesized by Integrated DNA Technologies (IDT) SiRNAs. At least two different siRNAs were used to target the same gene. The sequences of these siRNAs are listed in Online Table III. Adenoviruses were generated and titrated by VectorBuilder (APEX-MOM, HA-RBX2, GFP, HA-OMP25) or made by ourselves (HA-Parkin). The expression of all these genes is driven by the Cytomegalovirus (CMV) promoter.

SH-SY5Y cells (CRL-2266) were cultured in MEM/F12 supplemented with 10% fetal bovine serum (FBS), 100 µg/mL of penicillin, 100 µg/mL of streptomycin in 5% CO2 at 37°C. HEK293FT cells (ThermoFisher, Cat# R70007) were grown in DMEM supplemented with 10% FBS at 37°C and 5% $CO_2$.

To create RBX2-knockout SH-SY5Y and Hela cell lines using CRISPR-Cas9 system, two different guide RNAs against Rbx2 (gRNA-1: 5'-GCGATACGTGCGCCATCTGC-3'; gRNA-2: 5'- CCACCGCGTTCCACTTCTTG-3') were cloned into plentiCRISPR v2 (Addgene, Cat# 49535) by GeneScript. Then, the recombinant lentiCRISPR plasmid was co-transfected with pVSVg (Addgene, Cat# 8454) and psPAX2 (Addgene, Cat# 12260) into HEK293FT cells using PolyJet™ In Vitro DNA Transfection Reagent (SignaGen Laboratories, Cat# SL100688) following the manufacturer's protocol for lentivirus production. SH-SY5Y and Hela ells were then infected by lentivirus with 10 µg/ml polybrene for 48 hours and followed by selection with 2 µg/ml puromycin (Sigma, Cat# P8833).

Some of the cells were treated with different compounds. The doses were described in either the figure legends or in the main text. The sources are listed in Online Table III.

#### Transmission Electron Microscopy (TEM)

TEM sample preparation and imaging was performed at the Electron Microscopy and Histology Core at Augusta University. Briefly, left ventricle apexes or cultured cardiomyocytes were fixed in 3.5% glutaraldehyde/Sorensen's phosphate buffer at room temperature for 24 h. The samples were dehydrated, ultrathin-sectioned using Leica EM UC6 Ultramicrotome and collected in 200-mesh copper grids. The sections were counterstained with 2% uranyl acetate in water from 20 minutes followed by a lead citrate solution and examined with JEOL JEM1400 Flash electron microscope (JEOL). Immunogold labeling with anti-HA antibodies was performed as previously reported[4].

**Proteolysis by proteinase K**

Fresh isolated mitochondria were incubated with various concentrations of proteinase K (RPI) in ice-cold HES buffer for 30 min on ice. The digestion was stopped by the addition of an equal volume of 2X SDS-PAGE sampling buffer. The samples were boiled for 10 min and centrifuged at 12,000 g for 10 min, the supernatant fractions were separated by SDS/PAGE and proteins were detected by Western blot analysis.

**Mitochondrial staining**

To assess mitochondrial membrane potential, cells were stained with 1 µM Tetramethylrhodamine (TMRM) (Life Technologies) and 1 µM Mito Tracker Green (Life Technologies) for 30 minutes. To measure the levels of mitochondrial reactive oxygen species, cells were stained with 5 µM MitoSox (Life Technologies) for 40 minutes. After the staining, cells were immediately washed with DMEM and confocal images were taken at 488 nm and 568 nm using confocal microscope (STELLARIS 8, Leica Microsystems). Over 100 cells per group were analyzed.

**Seahorse XF assay**

The oxygen consumption rate (OCR) was determined by Seahorse Bioscience XFe24 Extracellular Flux Analyzer (Agilent Technologies) using XF Cell Mito Stress Test kit (Agilent Technologies) according to the manufacturer's instructions. Briefly, rat neonatal cardiomyocytes were seeded in Seahorse XF-24 plates at 50,000 cells/well in media containing 10% FBS for 48 hours. Cells were then transfected with siRNAs for another 48 hours. On the day of the analysis, the microplate was incubated in XF mitochondrial stress media (XF base medium add 10 mM Glucose, 2mM Glutamine, 1 mM Pyruvate) at 37°C for 1 hour without $CO_2$ before measurement. After OCR was recorded at baseline, oligomycin (2 µM), FCCP (1 µM) and rotenone/antimycin A (0.5 µM) were sequentially added to reveal the key parameters showing metabolic function. Finally, basal respiration, ATP production, maximal respiration, spare capacity and proton leak were calculated. All measurements were normalized to total proteins. Mitochondrial OCR was calculated by subtracting the final OCR value (pmol $O_2$/min) after rotenone/antimycin A treatment from the average of the three OCR values before rotenone/antimycin A treatment.

**Isolation of mitochondria**

To isolate mitochondria from NRVCs, the cells were washed once in PBS and homogenized in ice-cold HES buffer (250 mM sucrose, 5 mM HEPES, 1mM pH 7.4 EDTA) using 20 forceful strokes of a pre-chilled tight glass pestle on ice. The homogenates were centrifuged twice at 500 g for 5 minutes. The supernatants were combined and centrifuged at 9000 g for 15 minutes. The mitochondrial pellets were washed three times in PBS before being subject to subsequent analyses.

##### Adeno-associated virus serotype 9 (AAV9) transduction

AAV9 expressing mt-Keima [5] under the control of the CAG promoter (AAV-mt-Keima) was created by Vigene Biosciences. Neonatal mice at the age of postnatal day 2 or 3 were subcutaneously injected with AAV-mt-Keima (1 x $10^{11}$ GC/pup). Genotyping of mouse pups was blind to injection performer.

##### RNA isolation and quantitative real-time PCR

Total RNA from NRVCs and mouse tissues were prepared using TRIzol (Thermo Fisher Scientific) according to the manufacturer's instructions and then reversely transcribed to cDNA with RevertAid RT Reverse Transcription Kit (Thermo Fisher Scientific). For quantitative real-time PCR analysis, an equal amount of cDNA was mixed with Power SYBR Green PCR master mix (Thermo Fisher Scientific) and real-time PCR was performed using a QuantStudio 5 Real-Time PCR System (Applied Biosystems). All primers used for qPCR are listed in Online Table III. The $2^{\Delta\Delta Ct}$ method was used to analyze the relative changes in gene expression normalized against rat *Rplp0* or mouse *Hprt* expression.

##### Western blot

Cardiac tissues or cultured cells were homogenized with 1% SDS lysis buffer (1% SDS, 50 Mm Tris pH8.0, 10 mM EDTA pH8.0) supplemented with protease and phosphatase inhibitors (Sigma-Aldrich). The extracts were used for Western blotting as described previously[2]. The primary antibodies used in this study are listed in the Online Table III. The blots were visualized with home-made ECL reagents [100 mM Tris-HCl pH 8.5, 1.24 mM luminol (Sigma, A8511), 0.196 mM p-Coumaric acid (Sigma, C9008), 0.009% $H_2O_2$]. Images were acquired with ChemiDoc MP Imaging system (Bio-Rad) and quantified with Image Lab software (Bio-Rad).

##### Echocardiography

Echocardiography was performed to record the cardiac function as we described before [2]. Briefly, mice were anesthetized by inhalation of isoflurane (2.5% for induction and 1.5% for maintenance) via a nose cone. Cardiac images and loops were recorded from short-axis view with visualization of both papillary muscles using a VEVO 2100 echocardiography system with MS400 (18-38 MHz) transducer (Visual Sonics). The LV echocardiographic morphometric and functional parameters, including EF%, FS%, IVS, LVPW, LVID, LV mass and heart rate were measured offline from the obtained M-mode images using VEVO 2100 software.

##### Histology and immunostaining

Hearts were excised, washed with saline solution, fixed in 4% paraformaldehyde, and embedded by either paraffin or O.C.T Compound (Tissue Tek). Tissue blocks were sectioned transversely close to the root of the papillary muscle to visualize the left and right ventricles. Myocardium sections (5-7 μm thick) were prepared and stained with hematoxylin and eosin (H&E) for histopathology or Fast Green (Sigma, Cat# F7252) and Direct Red 80 (Sigma, Cat# 365548) staining and imaged by light microscopy.

For immunostaining, cardiac slices or neonatal cardiomyocytes were permeabilized with 0.1% Triton X-100 in PBS for 15 minutes, blocked with 10% goat serum blocking buffer for 1 hour, immunostained with primary antibodies overnight at 4°C, and subsequently incubated with appropriate Alexa-Fluor conjugated secondary antibodies (Thermo Fisher Scientific) for 1 h at room temperature. Finally, sections or slides were stained with DAPI (Sigma) and mounted in VECTASHIELD antifade mounting medium (Vector Laboratories). Images were captured with Zeiss Upright 780 confocal microscope (Zeiss) or STELLARS Confocal microscope (Leica Microsystems). The acquired images were quantified by ImageJ software (NIH). Section or slide information was pre-coded and blind to procedure performers including embedding, sectioning, staining, imaging and quantification. Antibodies used for immunostaining include HA (Sigma, Cat# H9658), TOMM20 (Cell Signaling Technology, Cat# 42406) Wheat germ agglutinin (Thermo Fisher Scientific, Cat# W11261), TUNEL (In Situ Cell Death Detection Kit, TMR red, Sigma, Cat#121567929), pUb (Cell Signaling Technology, Cat# 70973), and HSP60 (Thermo Fisher Scientific, Cat# MA3-012).

**Isolation of adult cardiomyocytes**

Adult cardiomyocytes were isolated by following an optimized Langendorff-free protocol [8]. Briefly, adult mice were anesthetized, and the heart was immediately flushed by injection of 7 mL EDTA buffer (concentration) into the right ventricle. The ascending aorta was clamped, and the heart was placed into a 60-mm dish containing fresh EDTA buffer. Digestion was achieved by sequential injection of 10 mL EDTA buffer, 3 mL perfusion buffer, and 30 to 50 mL collagenase buffer containing 0.5 mg/ml Collagenase 2 (Worthington Biochemical, Cat# LS004176) 0.5 mg/ml Collagenase 4 (Worthington Biochemical, Cat# LS004188), 0.05 mg/ml Protease XIV in perfusion buffer into the left ventricle through the same hole. Tissues were then minced into 1-mm pieces using forceps and gently triturated with a wide-bore 1 mL pipette. Collagenase enzyme activity was inhibited by addition of 5 mL stop buffer (5% FBS in Perfusion buffer). The cell suspension was passed through a 100-µm filter, and cells underwent 3 sequential rounds of gravity settling, using 3 intermediate calcium reintroduction buffers (0.34, 0.68 and 1.02 mmol/l $Ca^{2+}$ respectively) to gradually restore calcium concentration to physiological levels. The yield and the percentage of viable rod-shaped cells were quantified using a hemocytometer. Cardiomyocytes were then pre-plated onto laminin (5 µg/ml, Sigma, Cat# L2020-1MG) precoated dishes in plating media containing 5% FBS in M199 media with 10 mM BDM (Sigma, Cat# B0753) for 3 hours and further cultured in culture media containing 0.1% BSA (Sigma, Cat# A8531), ITS (Gibco, Cat# 41400045 ), CD lipid (Gibco, Cat# 11905031) and 25 µM blebbistatin (Sigma, Cat# 203390) in M199 media [9].

**Cell death and cytotoxicity analysis**

Live and dead cell staining was performed using the LIVE/DEAD Viability/Cytotoxicity Kit for mammalian cells (Thermo Fisher Scientific, L3224) according to the manufacturer’s instructions. Briefly, cells were incubated with Hoechst 33342 (10 μg/ml), calcein (1 μM) and ethidium (0.5 μM) dyes in M199 (Thermo Scientific) culture medium for 30 min at 37°C to identify nucleus, live and dead cells, respectively. The cells were then washed gently with DPBS and imaged using standard fluorescence microscopy (Nikon Eclipse Ti).

Lactate dehydrogenase activity (LDH) in medium, indicative of cytotoxicity, was measured using Cytotoxicity Detection Kit (Roche, Cat# 11644793001) by following the manufacturer's instructions.

**Statistics**

Statistical analyses were performed with GraphPad Prism. All values are presented as mean±SEM. Two-tailed parametric *t* tests were used for 2-sample comparisons if the values were normally distributed. Otherwise, the Mann-Whitney U test was used. A one-way analysis of variance with a Tukey post hoc test was used for comparisons among multiple groups. A Kaplan-Meier survival plot was deployed to evaluate mouse survival and Log-rank test was used to calculate statistical differences. Values of $P<0.05$ were considered statistically significant.
