## Supplemental Table 3 for "The Ubiquitin Ligase RBX2/SAG Regulates Mitochondrial Ubiquitination and Mitophagy"

**Online Table 3. Oligonucleotides, antibodies and compounds**

**Adenovirus**

| **Name** | **Vendor** | **Cat#** |
| --- | --- | --- |
| Ad-APEX2-MOM | Vectorbuilder | AVS(VB210419-1169guy)-C |
| Ad-HA-RBX2 | Vectorbuilder | AVM(VB210129-1063yxt)-C |
| Ad-GFP | Vectorbuilder | AVM(VB150925-10024)-C |
| Ad-HA-OMP25 | Vectorbuilder | AVM(VB210417-1068qst)-C |
| Ad-HA-parkin |  | Homemade |
| Ad-PINK1 |  | Homemade |

**Compound**

| **Name** | **Vendor** | **Cat#** |
| --- | --- | --- |
| Carbonyl cyanide m-chlorophenyl hydrazon (CCCP) | Sigma | C2759 |
| proteinase K | Thermo Fisher Scientific | AM2542 |
| E64d | Sigma | E8640 |
| Pepstatin A | Sigma | P5318 |
| BFA | Sigma | B1793 |
| FCCP | Sigma | C2920 |
| Oligomycin | Sigma | O4876 |
| Rotenone | Sigma | 557368-1GM |
| Antimycin A | Sigma | A8674 |
| Brdu | Sigma | B5002 |
| Tamoxifen | Sigma | T5648 |
| Biotin-phenol | AdipoGen Life Sciences | CDX-B0270 |
| Biotin | Sigma | B4501-100MG |
| (S)-6-Hydroxy-2,5,7,8-tetramethylchromane-2-carboxylic acid (Trolox) | TCI America | 53188-07-1 |
| Sodium azide | Sigma | S8032-25G |
| Sodium ascorbate | Sigma | 11140-50G |
| Hydrogen peroxide solution | Sigma | H1009-5ml |

**Genotyping primers**:

| **Name** | **Sequences** |
| --- | --- |
| αMHC-Cre-F | ATGACAGACAGATCCCTCCTATCTCC |
| αMHC-Cre-R | CTCATCACTCGTTGCATCATCGAC |
| RBX2-flox-F | TTCTGGCCAGGTGTGGTGATATC |
| RBX2-flox-R | CTTAGCCTTGGTTGTGTAGAC |
| MCM-F | CGTCCTCCTGCTGGTATAG |
| MCM-R | GTCTGACTAGGTGTCCTTCT |
| Des J | CAGCTTCAGGAACAGCAGGTCC |
| Des K | CATCAATCTCGCAGGTGTAGGACT |
| oIMR7026 PRKN-1 | CCTACACAGAACTGTGACCTGG |
| oIMR7027 PRKN-2 | GCAGAATTACAGCAGTTACCTGG |
| oIMR7028 PRKN-3 | ATGTTGCCGTCCTCCTTGAAGTCG |

**Q-PCR Primers**

| **Gene**  **Name** | **Species** | **Sense** | **Antisense** |
| --- | --- | --- | --- |
| *Nppa* | mouse | CACAGATCTGATGGATTTCAAGA | CCTCATCTTCTACCGGCATC |
| *Nppb* | mouse | GTCAGTCGTTTGGGCTGTAAC | AGACCCAGGCAGAGTCAGAA |
| *Myh6* | mouse | GGGCTGGAGCACTGAGAG | GAGAGAGGAACAGGCAGGAA |
| *Myh7* | mouse | CGCATCAAGGAGCTCACC | CTGCAGCCGCAGTAGGTT |
| *Acta1* | mouse | AATGAGCGTTTCCGTTGC | ATCCCCGCAGACTCCATAC |
| *Atp2a2* | mouse | TCGACCAGTCAATTCTTACAGG | CAGGGACAGGGTCAGTATGC |
| *Rbx2* | rat | CGTCCTTTCTTCGCACTCC | GGCTACCGCGTTCCACTTC |
| *Col1a* | rat | AGACCTGTGTGTTCCCTACT | GAATCCATCGGTCATGCTCTC |
| *Col3a* | rat | GGCTGCAAGATGGATGCTATAA | GAATCTGTCCACCAGTGCTTAC |
| *HPRT* | rat | GGCCAGACTTTGTTGGATTTG | CGCTCATCTTAGGCTTTGTATTTG |
| *Rbx2* | rat | GGTGATGGATGCCTGTCTTAG | GTTGTGGAAGGAGTGGTTACA |
| *Tomm20* | rat | GCTGAGGATGATGTGGAATGA | CAAGCACAGTTTGCCCTTATC |
| *Vdac1* | rat | GGAGTTTGGTGGCTCCATTTA | GACCTGATACTTGGCTGCTATTC |
| *Cisd1* | rat | GGCCTTGCCATTGAAACATC | GGATCCTTACGAGACCAAGATAAC |
| *Rhot2* | rat | CCATCATCCTGGTAGGCAATAA | ACTCCACACAGGTCTCTATCT |
| *Samm50* | rat | CACTCATTGAAGTCGTCTCTCTC | CAGCTCCTGGTTGACTTTGA |
| *Acsl1* | rat | AAGCTTGCAGGCCTTTCT | GGACCCTTGAAATCCTCTCTTC |
| *Hsp60* | rat | CTGTTCTGGCACGGTCTATT | GCATCAACAGCCAACATCAC |
| *Chchd3* | rat | GTGTGAGCCTCTTGCTATGT | TCTCTCCTGTCTGTCTGTCTT |
| *Agk* | rat | TGACGAGCAAAGAGGACTTTAT | AGGGTCTCTCACCTTCTTACT |
| *Cyc1* | rat | ATTGCGAGAAGGCCTCTATTT | GTACCATCGTCATACTCCAAGAC |
| *Ndufs5* | rat | GCATTAGCCTAGATCGACACTTTA | TCCCATGTGCGCATTCTATC |
| *Sod2* | rat | TAGAGCCTTTGCCTGTCTTATG | CAATGTCACTCCTCTCCGAATTA |
| *Rplp0* | rat | TTGAAATCCTGAGCGATGTGCAGC | GCCATTGTCAAACACCTGCTGGAT |

**SiRNAs from IDT**

| **Name** | **Target sequence** |
| --- | --- |
| Rat siRBX2-b | 5’- AAUGUAACCACUCCUUCCACAACTG-3’ |
| Rat siRBX2-c | 5’- GGAUGCCUGUCUUAGAUGUCAAGCT-3’ |
| Rat siParkin-a | 5’-GAAUCACCUGACAGUACAGAACUGT-3’ |
| Rat siParkin-b | 5’-GGGAUGAUGUCUUAAUUCCAAACCG-3’ |
| Mouse siRBX2-b | 5’-GUAGUCCAAAGAAUCGGCAAAUGAG-3’ |
| Mouse siRBX2-c | 5’-GUAGUCCAAAGAAUCGGCAAAUGAG-3’ |
| siLuciferase | 5’-AACGUACGCGGAAUACUUCGA-3’ |

**Primary antibodies**

| **Proteins** | **Vendor** | **Cat#** | **Dilution** |
| --- | --- | --- | --- |
| RBX2 | Abcam | ab181986 | 1:1000 |
| CUL5 | Bethyl | A302-173A-T | 1:1000 |
| RBX1 | Cell Signaling Technology | 11922s | 1:1000 |
| NAE1 | Cell Signaling Technology | 14321 | 1:1000 |
| UBC12 | Epitomics | 3690-1 | 1:1000 |
| UBE1 | Novus Bio | NBP1-90307 | 1:1000 |
| VDAC | Invitrogen | PA1-954A | 1:1000 |
| TUBULIN | DSHB | E7 | 1:5000 |
| Biotin, HRP-linked | Cell Signaling Technology | 7075s | 1:1000 |
| TOMM20 | Cell Signaling Technology | 42406 | 1:1000 |
| SOD2 | Cell Signaling Technology | 13141 | 1:1000 |
| Total OXPHOS Rodent WB Antibody Cocktail | Abcam | ab110413 | 1:1000 |
| Phospho-Ubiquitin (Ser65) | Cell Signaling Technology | 62802S | 1:1000 |
| P62 | American Research Products | GP62-C | 1:5000 |
| Ub (VU-1) | LifeSensors | VU101 | 1:1000 |
| GAPDH | Sigma | G8795 | 1:2000 |
| LC3 | MBL International Corporation | M186-3 | 1:1000 |
| CUL3 | Novus Biologicals | NB100-58788 | 1:1000 |
| CUL4a | Novus Biologicals | NB100-2267 | 1:1000 |
| TOMM40 | Cell Signaling Technology | 55959 | 1:1000 |
| CISD1 | Proteintech | 16006-1-AP | 1:1000 |
| RHOT2 | Proteintech | 11237-1-AP | 1:1000 |
| SAMM50 | Abclonal | A3401 | 1:1000 |
| ACSL1 | Abclonal | A16253 | 1:1000 |
| HSP60 | Thermofisher Scientific | MA3-012 | 1:1000 |
| PINK1 | Novus Biologicals | BC100-494 | 1:1000 |
| PARKIN | Cell Signaling Technology | 4211S | 1:1000 |
| pPARKIN | Cell Signaling Technology | 36866S | 1:1000 |
| Cleaved Caspase3 | Cell Signaling Technology | 9664L | 1:1000 |
| CHCHD3 | Abclonal | A8584 | 1:1000 |
| AGK | Santa cruz | sc-514235 | 1:200 |
| CYC1 | Proteintech | 10242-1-AP | 1:1000 |
| NDUFS5 | Proteintech | 15224-1-AP | 1:1000 |
| HA | Cell Signaling Technology | 3724s | 1:1000 |
| TRAF2 | Santa cruz | sc-7346 | 1:1000 |
| PPEF2 | Invitrogen | PA5-48692 | 1:1000 |
| PGAM5 | Cell Signaling Technology | 24584S | 1:1000 |
| PTEN | Cell Signaling Technology | 9559S | 1:1000 |
| BNIP3L | Cell Signaling Technology | 12396S | 1:1000 |
| BNIP3 | Abcam | ab10433 | 1:1000 |
