## Supplemental Table 4-7 for "The Ubiquitin Ligase RBX2/SAG Regulates Mitochondrial Ubiquitination and Mitophagy"

**Online Table 4. Echocardiography of RBX2 iCKO and control mice before and after tamoxifen injection (50 mg/kg).**

| Weeks | Pre | | | after | | |
| --- | --- | --- | --- | --- | --- | --- |
|  | **F/F** | **MCM** | **iCKO** | **F/F** | **MCM** | **iCKO** |
| N | 17 | 6 | 23 | 12 | 6 | 12 |
| IVS;d (mm) | 0.84±0.04 | 0.98±0.02 | 0.82±0.03 | 0.82±0.04 | 0.92±0.04 | 0.68±0.04* |
| IVS;s (mm) | 1.4±0.05 | 1.6±0.05 | 1.43±0.03 | 1.49±0.06 | 1.35±0.07 | 0.86±0.05**** |
| LVID;d (mm) | 3.23±0.08 | 2.93±0.25 | 3.15±0.06 | 3.17±0.22 | 3.26±0.13 | 4.16±0.22*** |
| LVID;s (mm) | 1.69±0.07 | 1.77±0.22 | 1.54±0.07 | 1.64±0.11 | 1.85±0.23 | 3.53±0.28**** |
| LVPW;d (mm) | 0.85±0.02 | 0.89±0.04 | 0.85±0.03 | 0.78±0.03 | 0.9±0.05 | 0.72±0.02 |
| LVPW;s (mm) | 1.29±0.05 | 1.44±0.06 | 1.36±0.04 | 1.3±0.07 | 1.42±0.06 | 0.92±0.07**** |
| EF (%) | 80.1±1.6 | 82.3±2.2 | 82.3±1.6 | 80.9±1.8 | 71.1±6 | 33±5.8**** |
| FS (%) | 48.1±1.6 | 50.2±2.3 | 50.8±1.8 | 50±1.9 | 40.9±4.8 | 16.1±3.1**** |
| LVVd (μl) | 42.5±2.5 | 40±4.7 | 40.3±1.9 | 45.9±3.1 | 50±7.5 | 78.8±8.7** |
| LVVs (μl) | 8.7±1 | 7.4±1.5 | 7.5±1 | 9±1.1 | 16.3±5.5 | 57.5±9.8**** |
| Heart rate (bpm) | 614±15 | 566±18 | 613±14 | 565±18 | 557±20 | 526±29 |

Data are Mean±SEM. *P < 0.05, **P < 0.01, ***P < 0.001 versus F/F. IVS;d, interventricular septal end-diastole; IVS;s, interventricular septal end-systole; LVID;d, LV internal dimension at end-diastole; LVID;s, LV internal dimension at end-systole; LVPW;d, LV posterior wall thickness at end-diastole; LVPW;s, LV posterior wall thickness at end-systole; EF, ejection fraction; FS, fraction shortening; LVVd, LV volume at end-diastole; LVVs, LV volume at end-systole.

**Online Table 5. Echocardiography of RBX2 iCKO and control mice** **before and after tamoxifen injection (20 mg/kg).**

| Weeks | Pre | | | 4 | | | 10 | | |
| --- | --- | --- | --- | --- | --- | --- | --- | --- | --- |
|  | **F/F** | **MCM** | **iCKO** | **F/F** | **MCM** | **iCKO** | **F/F** | **MCM** | **iCKO** |
| N | 9 | 6 | 14 | 10 | 6 | 11 | 7 | 6 | 6 |
| IVS;d (mm) | 0.87±0.03 | 0.8±0.1 | 0.79±0.04 | 0.78±0.04 | 0.92±0.03 | 0.79±0.04 | 0.84±0.04 | 1.03±0.08 | 0.84±0.05 |
| IVS;s (mm) | 1.48±0.06 | 1.5±0.06 | 1.42±0.05 | 1.47±0.06 | 1.73±0.07 | 1.47±0.14 | 1.57±0.06 | 1.84±0.07* | 1.46±0.1 |
| LVID;d (mm) | 3.38±0.08 | 3.26±0.07 | 3.3±0.08 | 3.39±0.07 | 3.17±0.06 | 3.67±0.06* | 3.5±0.1 | 3.19±0.13 | 3.69±0.1 |
| LVID;s (mm) | 1.62±0.13 | 1.42±0.11 | 1.65±0.08 | 1.56±0.11 | 1.67±0.04 | 2.3±0.18** | 1.64±0.1 | 1.33±0.15 | 2.25±0.2* |
| LVPW;d (mm) | 0.86±0.03 | 0.76±0.02 | 0.82±0.04 | 0.75±0.03 | 0.92±0.01 | 0.93±0.11 | 0.77±0.03 | 0.89±0.02 | 0.81±0.05 |
| LVPW;s (mm) | 1.47±0.07 | 1.53±0.06 | 1.39±0.04 | 1.4±0.03 | 1.55±0.08 | 1.06±0.08* | 1.42±0.06 | 1.53±0.06 | 1.27±0.1 |
| EF (%) | 83.1±2.8 | 84.6±3.4 | 82.2±1.7 | 85.3±1.7 | 80.4±1.8 | 62.2±7.1** | 84.4±1.7 | 87.7±3.3 | 69.5±4.9* |
| FS (%) | 52.3±3.5 | 52.9±3.9 | 50.2±1.6 | 54.2±2.2 | 48±2.1 | 39±4.5** | 52.9±2.2 | 57±4.6 | 39.5±4.5* |
| LVVd (μl) | 46.3±2.8 | 42±2.8 | 44.8±2.5 | 46.7±2.6 | 45.5±7.1* | 61.8±5.7* | 50.6±3.9 | 45±6.7 | 58±3.8 |
| LVVs (μl) | 8.1±1.4 | 6.5±1.4 | 8.3±1.1 | 7.2±1.1 | 8.7±0.7 | 26.4±8.2** | 7.9±1.2 | 6.3±2.3 | 18.2±3.8* |
| Heart rate (bpm) | 586±20 | 598±8.5 | 587±9.7 | 642±18 | 693±45 | 628±15 | 688±32 | 631±32 | 656±29 |

Data are Mean±SEM. *P < 0.05, **P < 0.01, ***P < 0.001 versus F/F. IVS;d, interventricular septal end-diastole; IVS;s, interventricular septal end-systole; LVID;d, LV internal dimension at end-diastole; LVID;s, LV internal dimension at end-systole; LVPW;d, LV posterior wall thickness at end-diastole; LVPW;s, LV posterior wall thickness at end-systole; EF, ejection fraction; FS, fraction shortening; LVVd, LV volume at end-diastole; LVVs, LV volume at end-systole.

**Online Table 6. Echocardiography of RBX2 CKO and control mice**

| Months | 1 | | | | 2 | | | 5 | | | | | 8 | | |
| --- | --- | --- | --- | --- | --- | --- | --- | --- | --- | --- | --- | --- | --- | --- | --- |
|  | **F/F** | α**MHC^Cre^** | **CKO** | **F/F** | | α**MHC^Cre^** | **CKO** | | **F/F** | α**MHC^Cre^** | **CKO** | **F/F** | | α**MHC^Cre^** | **CKO** |
| N | 9 | 6 | 15 | 9 | | 7 | 16 | | 7 | 11 | 11 | 6 | | 12 | 10 |
| LVID;d (mm) | 3.11±0.09 | 3.32±0.09 | 3.33±0.06 | 1.67±0.08 | | 1.79±0.15 | 1.83±0.06 | | 3.1±0.09 | 3.15±0.1 | 3.19±0.1 | 3.2±0.21 | | 3.89±0.18** | 3.94±0.16** |
| LVID;s (mm) | 1.77±0.11 | 2.05±0.15 | 2.11±0.08** | 1.67±0.08 | | 1.79±0.15 | 1.83±0.06** | | 1.67±0.02 | 1.84±0.08 | 2.0±0.07* | 1.67±0.16 | | 2.84±0.12** | 3.22±0.25** |
| LVPW;d (mm) | 0.67±0.03 | 0.7±0.05 | 0.62±0.04 | 0.64±0.03 | | 0.69±0.04 | 0.67±0.02 | | 0.68±0.07 | 0.67±0.03 | 0.67±0.03 | 0.93±0.06 | | 0.82±0.03 | 0.74±0.05** |
| LVPW;s (mm) | 1.05±0.04 | 1.07±0.08 | 0.96±0.04 | 1.1±0.06 | | 1.17±0.05 | 1.06±0.03 | | 1.2±0.05 | 1.19±0.02 | 1.04±0.04* | 1.32±0.05 | | 1.2±0.03 | 0.8±0.07*** |
| EF  (%) | 73.9±2 | 72.2±2.6 | 65.9±1.7* | 78.1±1.9 | | 78.4±2.5 | 71.6±1* | | 77.5±1.5 | 73.4±1.4 | 69±1*** | 80.2±3 | | 54.4±2.9*** | 36±4.2**** |
| FS  (%) | 41.8±1.8 | 40.6±2.3 | 35.5±1.2* | 45.7±1.8 | | 46.2±2.3 | 40.23±1.1* | | 44.41±1.9 | 41.5±1.2 | 37.1±0.6*** | 48.3±3.1 | | 28.1±1.9**** | 17.3±2.3**** |
| LVVd  (μl) | 38±5.1 | 41.9±0.9 | 44.8±2.1 | 33.3±3.9 | | 42.1±4.9 | 37±2.3 | | 33.3±4.6 | 40.1±2.8 | 41.5±3 | 37.2±7.2 | | 68.7±5.3** | 66.1±7.4* |
| LVVs  (μl) | 12.3±2.8 | 11.6±1.2 | 14.5±1.4 | 8.2±1 | | 10.5±2.2 | 10.4±0.9 | | 8.2±0.2 | 10.5±0.8* | 13±1.2** | 8.7±2 | | 31.6±3.4*** | 43.6±7.9*** |
| Heart rate (bpm) | 562±19 | 583±15 | 574±13 | 597 ±27 | | 604±21 | 621±10 | | 617±25 | 623±15 | 598±7 | 695±39 | | 610±16 | 632±31 |

Data are Mean±SEM. *P < 0.05, **P < 0.01, ***P < 0.001 versus F/F. LVID;d, LV internal dimension at end-diastole; LVID;s, LV internal dimension at end-systole; LVPW;d, LV posterior wall thickness at end-diastole; LVPW;s, LV posterior wall thickness at end-systole; EF, ejection fraction; FS, fraction shortening; LVVd, LV volume at end-diastole; LVVs, LV volume at end-systole.

**Online Table 7. Echocardiography of Parkin and RBX2 double knockout mice**

| **Months** | **5** | | | | **8** | | | | |
| --- | --- | --- | --- | --- | --- | --- | --- | --- | --- |
|  | **Parkin^-/-^** | **RBX2^Het^/**  **Parkin^-/-^** | **RBX2^CKO^** | **RBX2^CKO^/Parkin^-/-^** | | **Parkin^-/-^** | **RBX2^Het^/**  **Parkin^-/-^** | **RBX2^CKO^** | **RBX2^CKO^/Parkin^-/-^** |
| **N** | 5 | 7 | 8 | 8 | | 7 | 7 | 8 | 7 |
| **IVS;d (mm)** | 0.73±0.02 | 2.23±0.55* | 0.82±0.04 | 0.73±0.03 | | 1.73±0.15 | 2.62±0.67 | 4.08±0.37**** | 2.24±0.59 |
| **IVS;s (mm)** | 1.32±0.06 | 1.68±0.2 | 1.34±0.07 | 1.3±0.05 | | 3.11±0.26 | 2.04±0.34 | 4.83±0.27 | 2.92±0.68 |
| **LVID;d (mm)** | 3.64±0.22 | 3.4±0.08 | 3.64±0.06 | 3.5±0.14 | | 3.45±0.08 | 3.95±0.07**** | 4.87±0.29**** | 4.3±0.06**** |
| **LVID;s (mm)** | 2.25±0.17 | 2.06±0.06 | 2.29±0.09 | 2.21±0.14 | | 1.85±0.05 | 2.9±0.1**** | 4.04±0.35**** | 3.39±0.15**** |
| **LVPW;d (mm)** | 0.76±0.03 | 0.78±0.04 | 0.8±0.04 | 0.76±0.03 | | 0.96±0.09 | 0.73±0.04* | 0.69±0.04** | 0.73±0.04* |
| **LVPW;s (mm)** | 1.12±0.05 | 1.23±0.03 | 1.19±0.04 | 1.22±0.05 | | 1.34±0.08 | 1.05±0.05* | 0.9±0.06*** | 0.97±0.07** |
| **EF (%)** | 78.6±3.5 | 73.2±2.9 | 73.8±0.9 | 74.7±2.3 | | 77.3±1.8 | 59.7±2.3**** | 33.9±5.7**** | 42.7±3.8**** |
| **FS (%)** | 47.1±3.6 | 41.8±2.5 | 42.2±0.8 | 43.1±2 | | 45.1±1.7 | 31.5±1.6**** | 16.6±3**** | 21±2.2**** |
| **LVVd (μl)** | 55.3±6.1 | 49.2±2.4 | 57.6±2.9 | 55.2±4.9 | | 46.2±1.6 | 71.6±2.9**** | 93.7±4.4**** | 80.6±3.3**** |
| **LVVs (μl)** | 12.5±2.7 | 13.1±1.4 | 15.3±1.2 | 14.6±2.4 | | 10.4±0.8 | 29.1±2.5**** | 68.2±10.6**** | 46.5±3.9**** |
| **Heart rate (bpm)** | 642±19 | 633±21 | 551±13 | 567±15 | | 639±28 | 556±9 | 633±19 | 562±27 |

Data are Mean±SEM. *P < 0.05, **P < 0.01, ***P < 0.001 versus Parkin^-/-^. IVS;d, interventricular septal end-diastole; IVS;s, interventricular septal end-systole; LVID;d, LV internal dimension at end-diastole; LVID;s, LV internal dimension at end-systole; LVPW;d, LV posterior wall thickness at end-diastole; LVPW;s, LV posterior wall thickness at end-systole; EF, ejection fraction; FS, fraction shortening; LVVd, LV volume at end-diastole; LVVs, LV volume at end-systole.
